# Mesodermal-niche interactions direct specification and differentiation of pancreatic endocrine cells in human multilineage organoids

**DOI:** 10.64898/2025.12.13.694117

**Authors:** Georgina Goss, Alejo Torres-Cano, Raquel A. Martinez García de la Torre, Siwanarta Ma, Leila Hermosilla Andrivet, Michelle Simon, Martina Pedna, Heather Wilson, Flavia Flaviani, Alessandra Vigilante, Francesca M. Spagnoli

**Affiliations:** Centre for Gene Therapy and Regenerative Medicine, King’s College London, Great Maze Pond, London SE1 9RT, UK; Hub for Applied Bioinformatics (HAB), King’s College London, Newcomen Street, London 17 SE1 1UL, UK

## Abstract

Tissues develop and function within complex microenvironments, where diverse cell types interact through tightly regulated spatial and temporal cues. These interactions shape cell identity, differentiation, and spatial organisation, ultimately establishing functional tissue architecture. Understanding and modelling these processes in humans therefore require multicellular systems that capture the cellular diversity and interactions of developing tissues. Here, we combine spatial transcriptomics of the human foetal pancreas with an induced pluripotent stem cell (iPSC)-based multilineage model that recapitulates key features of human endocrine pancreatic development and incorporates a mesoderm-derived cellular niche. We comprehensively characterise the multilineage organoid transcriptomes across differentiation stages and benchmark their fidelity against single-cell and spatial transcriptomic profiles of human foetal pancreas. We demonstrate that iPSC-derived pancreatic mesenchymal cells promote endocrine commitment by restricting off-target cell fates and enhancing β-cell differentiation. Together, these findings establish a framework for investigating the cellular mechanisms underlying human endocrine development and disease.

## Main

During pancreatic organogenesis, crosstalk between the pancreatic epithelium and its surrounding mesenchymal tissue orchestrates lineage allocation, morphogenesis and the emergence of functional endocrine cells^1–5^. Despite the recognised importance of the pancreatic mesenchyme, there is currently no consensus on the number or identity of distinct mesenchymal populations in the human pancreas, their precise spatial positioning relative to endocrine populations, how these niches shift across developmental time, or which subtypes provide the instructive cues that drive beta-cell differentiation and function^6–12^. This uncertainty underscores the need for integrated approaches that combine *in situ* mapping with functional co-culture systems to directly link mesenchymal identities to beta-cell differentiation/maturation potential. Prior studies have attempted to recreate aspects of the pancreatic mesenchymal niche by combining primary or immortalised fibroblasts or endothelial cells with stem cell-derived pancreatic progenitors or islet-like clusters, resulting in increased expression of insulin as well as improved glucose responsiveness^13–17^. Whilst these co-culture approaches can enhance certain beta-cell characteristics, they rely on components sourced from primary tissues that are neither tissue-specific nor stage-matched. As a result, they fall short to fully capture the endogenous intercellular communication that occurs during *in vivo* human pancreatic development. These fundamental limitations highlight the need for a human system to model islet development as well as to further understand disease mechanisms and therapeutic strategies.

To recapitulate the cellular diversity and interactions that occur *in vivo*, here we developed an approach that combines spatial transcriptomics of human foetal pancreas with an *in vitro* multilineage organoid system derived from human iPSCs. This strategy directly connects *in situ* niche signatures to the mesenchymal-epithelial interactions underlying pancreatic and endocrine cell fate specification and beta-cell differentiation.

### Dynamic spatial organization of the human pancreas during development

To map the cellular composition of the human pancreatic microenvironment and its spatial relationships to the major pancreatic epithelial cell types-ductal, acinar, and endocrine-, we performed CosMx™ Spatial Molecular Imaging (SMI), which allows visualization of transcript distribution at single-cell resolution. We analysed tissue sections collected from three stages of human pancreatic development, spanning 8-19 post-conception weeks (PCW), capturing the progression from pancreatic fate specification to differentiation and morphogenesis^9,18^ (Fig. 1 and Extended Data Fig. 1).

**Fig. 1.**
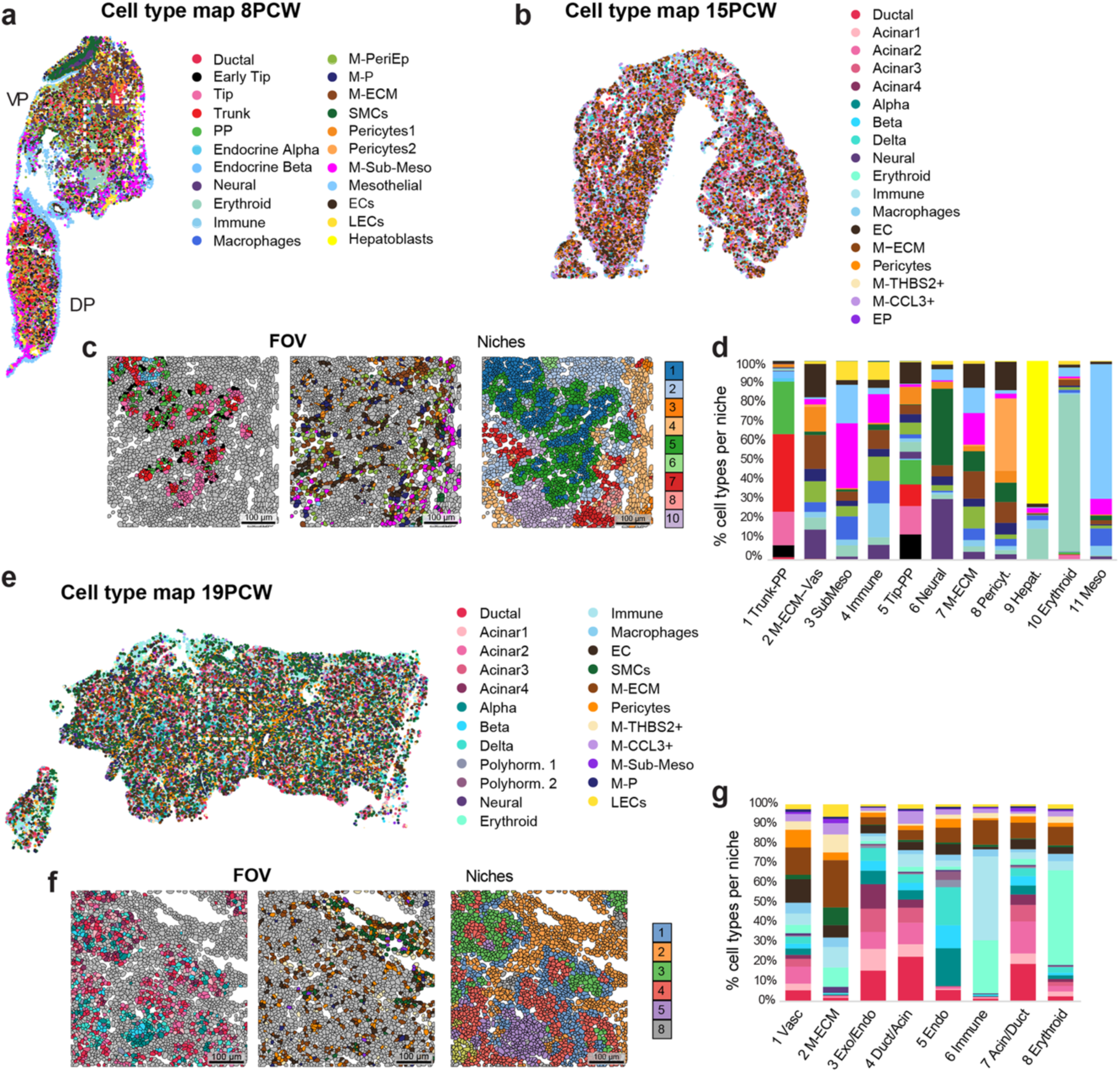
Single-cell spatial transcriptomics by CosMx SMI of human foetal pancreatic tissue. **a,** Spatial distribution of annotated cell populations in a representative 8 Post-Conception Weeks (PCW) pancreatic tissue section. Each colour represents a cell type. Dorsal (DP) and ventral (VP) pancreatic buds are indicated. White dashed box indicates a representative field of view (FOV), shown at high magnification in (c). **b**, Spatial distribution of annotated cell populations in a representative 15PCW pancreatic tissue section. Each colour represents a cell type. **c**, Representative fields of view (FOV) of an 8PCW tissue section highlighting pancreatic epithelial (left) or mesenchymal and vascular (middle) populations or niches (right). **d,** Proportion of cell types per BANKSY niche at 8PCW; cell types are coloured as in (a). **e,** Spatial distribution of annotated cell populations in a representative 19 PCW pancreatic tissue section. Each colour represents a cell type. White dashed box indicates a representative FOV, shown below at high magnification in (f). **f,** Representative FOV of a 19PCW tissue section, highlighting pancreatic epithelial (left) or mesenchymal and vascular (middle) cell populations or niches (right). **g,** Proportion of cell types per BANKSY niche at 19PCW; cell types are coloured as in (e).

Following cell segmentation and cell identification, the corresponding transcriptomic data were exported from the AtoMx platform and imported into Seurat. Using an analysis pipeline analogous to that applied to single-cell RNA-sequencing (sc-RNASeq) data, we generated UMAP embeddings for each sample and assigned cell-type identities to the resulting clusters (Fig. 1a,b,e and Extended Data Fig. 2a-f). Integration of our CosMx spatial data with published *in vivo* human pancreatic datasets^6,7^ showed robust overlap and lineage-specific correlations in both pancreatic epithelium and surrounding microenvironment compartments across developmental stages (Extended Data Fig. 3). Furthermore, we confirmed cell type annotations based on the expression of known marker genes present in the spatial panel (Extended Data Fig. 2g).

At 8PCW, the dorsal and ventral pancreatic buds were still distinguishable (Fig. 1a). Although both buds contained multipotent pancreatic progenitors (PP), early tip and tip progenitors, trunk progenitors and endocrine cells, differences in the abundance of specific populations were evident. For example, PP and Trunk progenitors were enriched in the dorsal bud, while large ductal progenitors were present in the ventral one (Fig. 1a and Extended Data Fig. 4a).

Based on tissue-specific marker genes and the top differentially expressed genes (DEGs), we annotated molecularly distinct cell clusters within the pancreatic microenvironment at 8PCW (Fig. 1a, Extended Data Fig. 2a,b and Supplementary Table 1), including neural and mesoderm-derived populations, such as immune, endothelial cells (ECs), as well as eight mesenchymal clusters, occupying distinct anatomical locations (Fig. 1a,c and Extended Data Fig. 4b). At the periphery of the pancreatic buds, we found the mesothelium (*MGP*^+^*, KRT19*^+^) cluster along with a sub-mesothelium population (characterized by elevated expression levels of *DUSP1* and *JUN*) and a *CCL21^+^*, *LYVE^+^* lymphatic endothelial (LEC) progenitor population (Extended Data Fig. 2b and Extended Data Fig. 4b). We also identified mesenchymal populations, which are typically associated with ECs (*PECAM1^+^*), such as pericytes (*PDGFRB^+^*) and smooth muscle cells (SMCs) (*ACTA2^+^*) (Extended Data Fig. 2b and Extended Data Fig. 4b). Finally, three mesenchymal populations were positioned closer to the epithelium: a population enriched for extracellular matrix (ECM) and signalling molecules (*e.g*., *BMP1*, *BMP4*, *WNT3*, *WNT5B*, *WNT5A*) [referred to as peri-epithelial (M-PeriEP)], a population characterized by the expression of proliferative genes (*MKI67*, *TOP2A*) (referred to as M-P) and a population with a gene signature associated with ECM components (M-ECM) (Extended Data Fig. 2b, Extended Data Fig. 4b and Supplementary Table 1). Notably, some of the mesenchymal populations, in particular the vascular-associated ones, were differentially enriched between the dorsal and ventral buds (Extended Data Fig. 4a,b).

At later stages (15PCW and 19PCW), tissue complexity increased both in cellular composition and structural organization. We identified acinar, ductal and endocrine cell types in more advanced states of differentiation, arranged in distinct acinar compartments and islet-like structures (Fig. 1b,e, Extended Data Fig. 2c-f and Supplementary Tables 2, 3). For example, we identified all major endocrine cell types, including beta, polyhormonal, alpha, and delta cells as well as four transcriptionally distinct acinar populations, which is consistent with previous reports^7,19,20^ (Fig. 1b,e). From 15PCW onward, we measured an overall decrease of the mesenchyme over the epithelial cell fraction, accompanied by changes in cellular composition, such as the emergence of distinct mesenchymal populations (M-*THBS2*^+^ and M-*CCL3*^+^ clusters) (Fig. 1b,e and Extended Data Fig. 2c-h).

Next, we identified spatial niches across samples by clustering and annotating cells according to their neighbourhood gene expression profiles using BANKSY^21^. At 8PCW, most niches were composed predominantly of mesoderm-derived populations, which is in line with the epithelial-mesenchymal ratio observed at this stage (Fig. 1c,d and Extended Data Fig. 2h). Despite the overall mesenchymal predominance, niches differed in their cellular composition. Some were enriched for M-Sub-Meso, LECs and immune populations, including macrophages or other immune cell progenitors (referred to as ‘Sub-Meso’ and ‘Immune’ niches), while another was characterized by an enrichment of M-ECM and vascular populations, including ECs and pericytes (‘M-ECM-Vas’) (Fig. 1d and Extended Data Fig. 4c,e). Pancreatic epithelial cells were organized in two main niches, comprising primarily tip (‘Tip-PP’) or trunk progenitors along with the first few specified endocrine cells (‘Trunk-PP’) (Fig. 1d and Extended Data Fig. 4e). The spatial arrangement of these two niches matched the expected tip-trunk organization of the embryonic pancreas, with the respective niches localized to the periphery and core, respectively. The ‘Tip-PP’ niche comprised a diverse range of mesenchymal populations, while the ‘Trunk-PP’ niche contained a smaller proportion of M-PeriEp and vascular supporting cells (Fig. 1d and Extended Data Fig. 4e). Neural progenitor cells co-localized predominantly with mesoderm-derived populations and were largely excluded from epithelial niches (Fig. 1d).

At 19PCW, the increased epithelial cellular fraction and greater cellular complexity were well reflected in the composition of ‘epithelial niches’, several of which were enriched in exocrine cell populations, while one was composed almost exclusively of endocrine cell types (‘Endo’) (Fig. 1f,g and Extended Data Fig. 4g,h). Notably, these epithelial niches also differed in their associated mesenchymal composition: the ‘Endo’ niche comprised preferentially vascular cells, while the ‘Duct/Acinar’ niche mainly comprised M-P and M-CCL3^+^ cell types (Fig. 1g and Extended Data Fig. 4h). The ‘Vascular’ and ‘M-ECM’ mesenchymal niches persisted at 19PCW, with both exhibiting a similar cellular composition as at earlier stages (Fig. 1g).

Together, our spatial and transcriptomic analyses revealed a highly organized pancreatic microenvironment, characterized by distinct mesenchymal populations that are closely coordinated with epithelial lineage differentiation. These findings establish a comprehensive cellular and spatial atlas of the developing human pancreas, providing a crucial framework for dissecting niche-driven mechanisms governing pancreatic organogenesis.

### Mesodermal-niche cues coax pancreatic differentiation towards the endocrine lineage in multilineage organoids

To recapitulate the *in vivo* cell-cell interactions and developmental niche dynamics, we aimed at establishing pancreatic organoids derived from human pluripotent stem cells, incorporating both endodermal and mesodermal components. First, to generate pancreatic-like mesodermal lineages, we adapted a previously established protocol^22–24^ to differentiate posterior foregut splanchnic mesoderm (referred to as pFG-SPM) from human iPSCs (Fig. 2 and Extended Data Fig. 5).

**Fig. 2.**
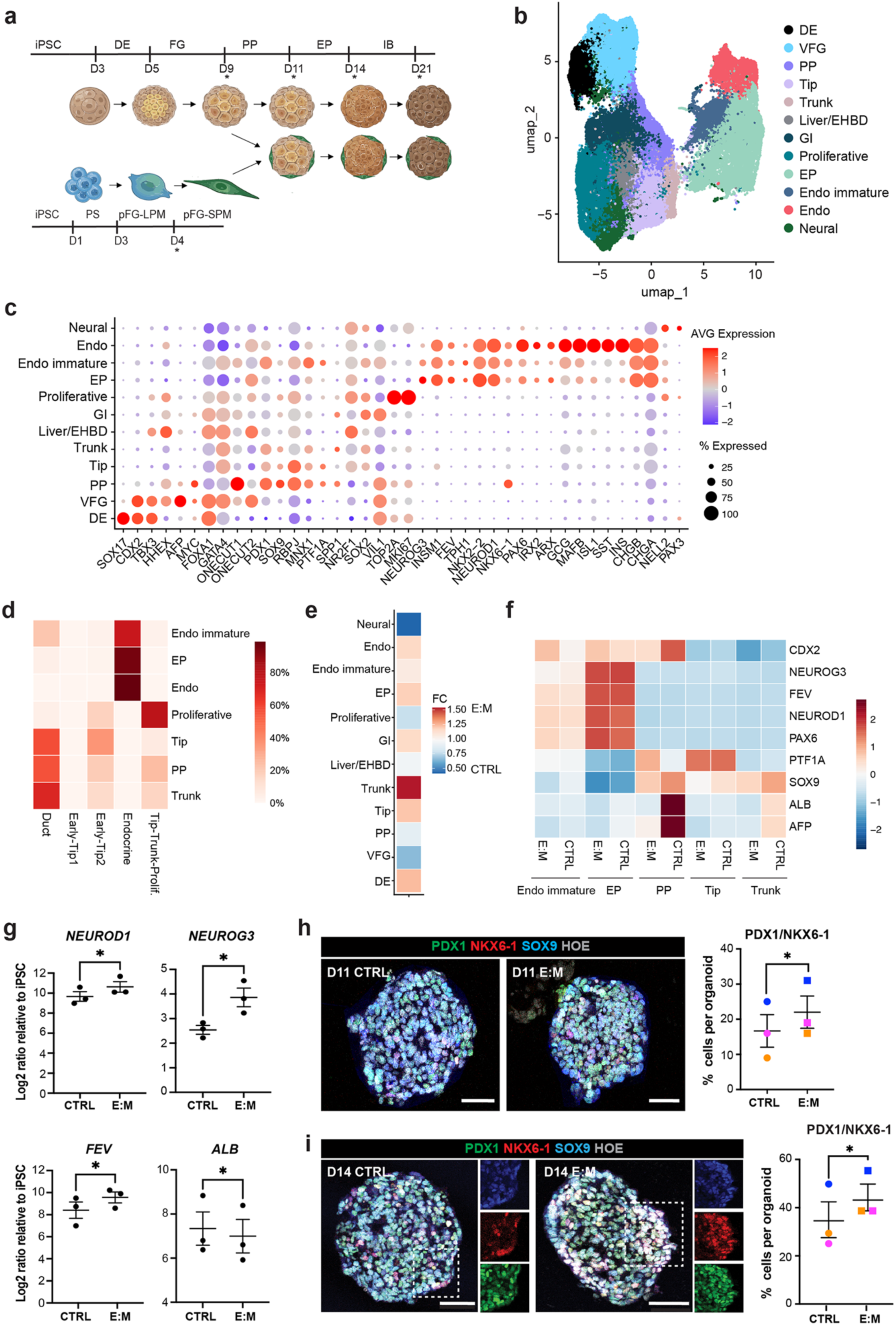
iPSC-derived pancreatic mesenchyme improves the fidelity of pancreatic progenitor differentiation. **a,** Schematic representation of iPSC-derived pancreatic progenitors (PP) and posterior foregut splanchnic mesoderm (pFG-SPM) co-culture strategy to establish multilineage organoids. Stars (*) indicate specific timepoints of the differentiation experiment chosen for further analysis by sc-RNASeq. Abbreviations: D, Day; DE, Definitive Endoderm; FG, Foregut; PP, Pancreatic Progenitor; EP, Endocrine Progenitor; IB, Immature Beta; PS, Primitive Streak; pFG-LPM, posterior Foregut Lateral Plate Mesoderm. **b**, UMAP visualization of the endoderm compartment from sc-RNASeq analysis on control (CTRL) and Epithelial:Mesenchyme (E:M) co-culture organoids. Abbreviations: DE, Definitive Endoderm; EHBD, Extra-Hepatic Biliary Duct; Endo, Endocrine cells; EP, Endocrine Progenitors; GI, Gastrointestinal; PP, Pancreatic Progenitors; VFG, Ventral Foregut. **c**, Dot Plot showing the expression of selected markers across populations identified. Colour represents scaled average (AVG) expression level; dot size indicates the percentage (%) of expressing cells (Supplementary Table S4). **d,** Co-occurrence heatmap showing the correspondence between pancreatic *in vitro* labels and transferred labels from *in vivo* dataset^6^ at equivalent developmental stages. **e,** Heatmap showing fold changes (FC) in cell-type abundance between E:M and CTRL organoid conditions. **f,** Mean scaled expression of indicated genes in endoderm clusters split per condition at D11 and D14. Colour scale represents scaled expression values of the indicated genes. **g,** RT-qPCR analysis of indicated endocrine and hepatic marker genes in CTRL and E:M organoids at D11. Data are shown as LOG2 ratio relative to undifferentiated iPSCs. Values shown are mean ± S.E.M. n=3. *p<0.05 (paired t-test). **h,** Left, representative immunofluorescence (IF) images of D11 CTRL and E:M organoids stained for PDX1 (green), NKX6-1 (red) and SOX9 (blue). Hoechst was used as nuclear counterstain (grey). Scale bar, 50µm. Right, plot showing the percentage (%) of PDX1^+^/NKX6-1^+^ cells relative to the total number of cells per organoid. Individual dots represent the mean value of biological replicates. Each biological replicate is colour-coded. Values shown are mean ± S.E.M. N= 3. (CTRL, n= 22; E:M, n= 35). *p<0.05 (paired t-test). **i,** Left, representative IF images of D14 CTRL and E:M organoids stained for PDX1 (green), NKX6-1 (red), SOX9 (blue), and Hoechst (grey). Right panels show boxed regions as single channels. Scale bar, 50µm. Right, plot showing the percentage (%) of PDX1^+^/NKX6-1^+^ cells relative to the total number of cells per organoid. Individual dots represent the mean value of biological replicates. Each biological replicate is colour-coded. Values shown are mean± S.E.M. N= 3. (CTRL, n= 27; E:M, n= 50). *p<0.05 (paired t-test).

Upon a four-day differentiation protocol, we obtained a heterogeneous mesodermal culture, including cells expressing pancreatic mesenchyme markers, such as *NKX2-5*, *ISL1*, *PBX1*, *FOXF1*, alongside cells positive for mesothelial and vascular marker genes, such as *TBX18, TCF21, WT1, ACTA2*, *RGS5* (Extended Data Fig. 5a,b). This heterogeneity aligns with previous reports, supporting that pancreatic mesenchymal populations derive from splanchnic mesenchyme^12,25,26^.

Next, we established a multilineage co-culture system by combining pFG-SPM with pancreatic endoderm lineage derived from an isogenic iPSC line (Fig. 2a). Briefly, iPSCs were differentiated to pancreatic progenitor (PP) stage [day (D) 9] using a previously established stepwise 3D suspension protocol^27^, mixed with a defined amount of pFG-SPM and cultured in a modified medium designed to preserve in culture both pancreatic endoderm and mesodermal lineages (Extended Data Fig. 5c; for details see Methods). When the two cell populations were combined in suspension, they aggregated in 3D organoids, with the epithelial component predominantly coalescing at the center and the mesenchymal cells remaining at the periphery (Extended Data Fig. 5d).

Both control (CTRL) and epithelial:mesenchyme (E:M) co-culture organoids were maintained in suspension and continuously characterized throughout the differentiation process by sampling organoids at multiple timepoints [day (D) 9, 11, 14 and 21] and performing sc-RNASeq using the 10x Genomics GEM FLEX-X platform, which enables the analysis of fixed samples (Fig. 2a and Extended Data Fig. 5e-g). As expected, unbiased clustering identified two main clusters corresponding to endodermal and mesodermal identities, along with a small fraction of ectodermal cells in the integrated organoid dataset. The mesodermal cluster was predominantly composed of cells from the starting pFG-SPM culture at D4, with one subset appearing specific to the E:M organoid culture, while the endodermal cluster was distributed across the different organoid populations (CTRL and E:M) and timepoints (Extended Data Fig. 5f).

We first focused on the endoderm compartment, representing the major cellular population, and examined transcriptional changes across differentiation stages and culture conditions. Clustering of the endoderm subset revealed 12 clusters distributed across the CTRL and E:M organoid datasets and different timepoints analysed (Fig. 2b and Supplementary Table S4). These clusters recapitulated the cellular diversity seen during foregut and pancreatic development^6^, encompassing multipotent pancreatic progenitors (PP) (*PDX1, SOX9, NKX6-1*), early tip (*PTF1A*) and trunk (*SPP1*) populations as well as a spectrum of endocrine populations, from endocrine progenitors (EP) to hormone-positive cells (Fig. 2c). A small number of clusters, predominantly present at the early timepoints (D11), represented early stages of foregut endoderm development or off-target populations, including other endodermal derivatives (GI and Liver/EHBD clusters) and cells with a neural-like signature (Fig. 2c). The transcriptional profiles of the identified clusters largely overlapped with sc-RNASeq datasets from *in vivo* foetal pancreatic tissue^6^ (Fig. 2d).

We next compared the cellular composition of the two conditions and observed a shift towards pancreatic cell identities in E:M organoids relative to CTRL condition. This was particularly evident in the enrichment of Trunk and Endocrine (EP and Endo) populations, accompanied by a reduction in off-target Liver/EHBD and neural-like clusters at D11 (Fig. 2e). Consistently, E:M organoids exhibited reduced expression of markers associated with off-target populations (*e.g*., *ALBUMIN*, *AFP*, *CDX2*) compared to CTRL at D11 and D14 (Fig. 2f,g). Changes in cellular composition were further corroborated by immunofluorescence (IF) assays, which showed a significant increase in PDX1/NKX6-1 double-positive (+) cells in E:M organoids (Fig. 2h,i). PDX1^+^/NKX6-1^+^ are considered bona-fide that can mature into functional, glucose-responsive insulin-producing beta-like cells^28,29^. Additionally, RT-qPCR analysis showed increased expression of genes critical for pancreatic endocrine development,^30–32^ such as *NEUROD1*, *NEUROG3* and *FEV* (Fig. 2g).

Consistent with these findings, density plot analysis of the endodermal transcriptomes at D11 revealed a shift in cell-state density in the E:M organoids, with high-density regions corresponding to Trunk and EP clusters (Extended Data Fig. 6a). In parallel, E:M organoids exhibited a reduced proportion of proliferative cells compared to CTRL, while apoptosis was extremely low in both conditions with non-significant differences (Extended Data Fig. 6b,c).

These results suggest that exposure to iPSC-derived pancreatic mesenchyme promotes progression towards more advanced stages of pancreatic differentiation.

Genes associated with acinar cell fate were only modestly induced and showed no significant differences between conditions (Extended Data Fig. 6d,e), suggesting that the established E:M conditions preferentially support endocrine lineage specification. Notably, endocrine differentiation was observed in E:M organoids, even when cultured in basal medium without exogenous pancreatic differentiation cytokines, reaching levels comparable to those observed in CTRL organoids (Extended Data Fig. 6f,g).

Together, these results demonstrate that integrating iPSC-derived PPs with pFG-SPM rapidly reshapes pancreatic lineage dynamics and enhances pancreatic lineage fidelity, promoting endocrine commitment, whilst limiting off-target differentiation.

### Multilineage iPSC-derived organoids are enriched in differentiated beta-like cells

To further characterize the endocrine differentiation potential of iPSC-derived E:M organoids, we targeted our transcriptome analysis to the endocrine compartment and complemented it with IF, RT–qPCR, and functional assays. Reclustering of endocrine cells from the integrated organoid dataset revealed 15 clusters, which were annotated based on DEGs and marker expression patterns defined in human foetal pancreas tissue^6,7,11^ (Fig. 3a,b and Supplementary Table S5). We identified all major islet cell types, including alpha (*GCG*, *IRX2*), beta (*INS*, *INSM1*), delta (*SST*, *HHEX*), as well as poly-hormonal cell states at D21. The remaining endocrine clusters corresponded to distinct endocrine progenitor states, characterized by the expression of endocrine progenitor transcription factors (*NEUROG3, NKX2-2, NEUROD1*), and enterochromaffin cells (*TPH1, CDX2, DDC*) (Fig. 3b). To further validate these annotations, we performed label transfer from a human foetal pancreatic sc-RNASeq atlas^6^ and our spatial 19PCW CosMx transcriptomics dataset, which yielded concordant assignments across major endocrine cell populations (Fig. 3c,d).

**Fig. 3.**
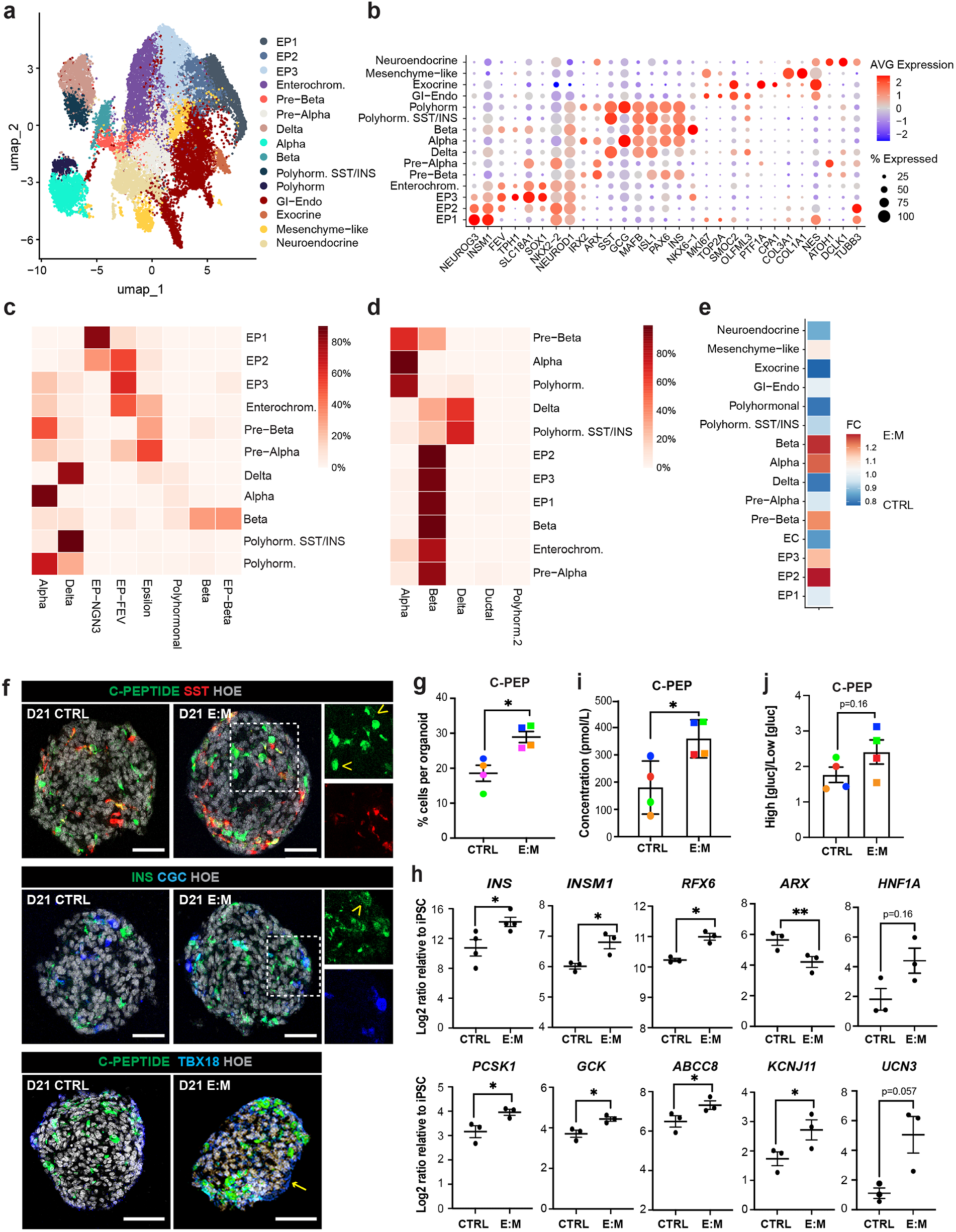
iPSC-derived pancreatic mesenchyme enhances beta-cell differentiation in multilineage organoids. **a**, UMAP visualization of the endocrine compartment from sc-RNASeq analysis on CTRL and E:M organoids. **b**, Dot Plot showing the expression of selected markers across populations identified. Colour represents scaled average (AVG) expression level; dot size indicates the percentage (%) of expressing cells (Supplementary Table S5). Abbreviations: EP, Endocrine Progenitors; Enterochrom., Enterochromaffin cells; GI-Endo, Gastrointestinal-Endocrine; Polyhorm., Polyhormonal cells. **c,** Co-occurrence heatmap showing the correspondence between endocrine *in vitro* labels and transferred labels from *in vivo* dataset^6^. **d**, Co-occurrence heatmap showing the correspondence between endocrine *in vitro* labels and transferred labels from the 19PCW CosMx dataset. **e**, Heatmap showing fold changes (FC) in cell-type abundance between E:M and CTRL organoid conditions. **f**, Representative IF images of D21 CTRL and E:M organoids stained for C-PEPTIDE (green), SOMATOSTATIN (SST) (red), and Hoechst (grey) (Top panel); INSULIN (INS) (green), GLUCAGON (GCG) (blue), and Hoechst (grey) (Middle panel); C-PEPTIDE (green), mesenchymal marker TBX18 (blue), and Hoechst (grey) (Bottom panel). Right panels show boxed regions as single channels. Arrowheads indicate monohormonal cells; arrows indicate TBX18^+^ mesenchymal cells at the outer layer of the organoid. Scale bar, 50μm. **g**, Quantification of the % of C-PEPTIDE^+^ cells relative to the total number of cells per organoid. Individual dots represent the mean value of biological replicates. Values shown are mean ± S.E.M. N= 4. (CTRL, n= 39; E:M, n= 66). *p<0.05 (paired t-test). **h,** RT-qPCR analysis of selected endocrine genes in CTRL and E:M organoids at D21. Data are shown as LOG2 ratio relative to undifferentiated iPSCs. Values shown are mean ± S.E.M. N=3-4. *p<0.05; **p=0.005 (paired t-test). **i**, ELISA assay measuring secreted C-PEPTIDE (pmol/L) in D21 CTRL and E:M organoid supernatants (N=4). Each biological replicate is colour-coded. Values shown are mean ± S.E.M. *p<0.05 (paired t-test). **j,** Representative static glucose stimulation insulin secretion (GSIS) assay of D21 CTRL and E:M organoids. On y-axis ratio of C-PEPTIDE secreted at high *versus* low glucose [gluc] conditions. Values shown are mean ± S.E.M. (N=4). *p<0.05 (paired t-test).

Endocrine clusters, especially the islet cell populations, were predominantly detected in late-stage organoid cultures (Extended Data Fig. 7a) and were enriched in E:M organoid conditions relative to CTRL. This enrichment was mostly pronounced in the pre-beta, beta and alpha populations (Fig. 3e).

We then performed IF analyses as orthogonal validation on both CTRL and E:M organoid cultures at D21. E:M organoids contained a higher proportion of monohormonal INSULIN^+^/C-PEPTIDE^+^ cells (Fig. 3f,g), consistent with enhanced beta-cell differentiation. This was accompanied by increased expression of *INS*, the beta-cell transcription factors, *INSM1*, *RFX6*, and genes involved in insulin processing and secretion, including *PCSK1*, *GCK*, *KCNJ11*, *ABCC8,* alongside reduced *ARX*, a transcription factor associated with alpha-cell fate^31^ (Fig. 3h and Extended Data Fig. 7b). Consistent with these changes, D21 E:M organoids exhibited increased C-Peptide secretion into the culture supernatant under basal conditions (Fig. 3i) and in response to glucose stimulation (Fig. 3j), although the latter increase did not reach statistical significance owing to variability between samples. Importantly, the enhanced endocrine differentiation observed in E:M organoids was accompanied by a reduction in the proportion of certain off-target cell populations, such as the enterochromaffin cells (Extended Data Fig. 7c-e).

To validate the robustness of our E:M multilineage organoid system, we used an additional iPSC line (AICS-0090), which showed comparable efficiencies in pFG-SPM differentiation and enhanced induction of pancreatic endocrine lineage (Extended Data Fig. 8).

### iPSC-derived mesenchyme promotes beta-cell differentiation through cell-cell interactions

To distinguish contact-mediated from secreted signalling, we supplemented CTRL pancreatic organoids with conditioned media (CM) from pFG-SPM between D9 to D20 of the culture and then analysed the different experimental conditions at the end of islet differentiation stage (Fig. 4a). The CM condition failed to recapitulate the enhanced differentiation seen in the E:M pancreatic organoids, where pFG-SPM was combined in direct contact with PPs. Specifically, D21 CM yielded fewer C-PEPTIDE^+^ cells compared to E:M organoids at the same stage and was indistinguishable from the mesenchyme-free CTRL organoids (Fig. 4b,c). Likewise, the expression of key beta-cell genes (*GCK, INSM1*, *RFX6*) was lower in D21 CM than in E:M D21 (Extended Data Fig. 9a). As expected, there was no induction of mesenchymal markers expression (Extended Data Fig. 9a) and the number of PDGFRB^+^ mesenchymal cells was comparable to those in D21 CTRL organoids (Fig. 4d,e).

**Fig. 4.**
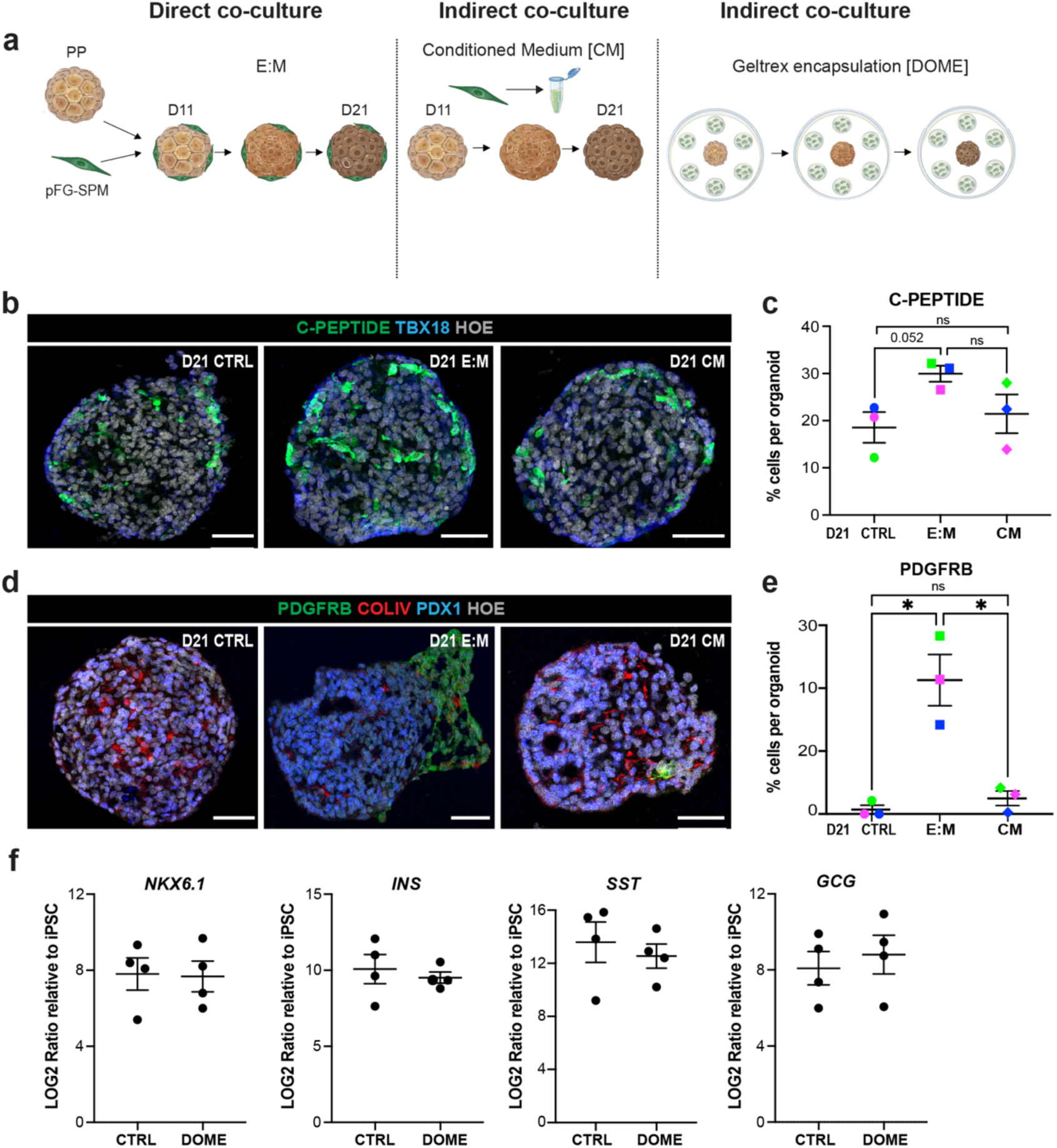
Cell contact-mediated cues enhance beta-cell differentiation in multilineage pancreatic organoids. **a,** Schematic representation of iPSC-derived pancreatic progenitors (PP) and posterior foregut splanchnic mesoderm (pFG-SPM) direct or indirect co-culture experimental conditions. **b,** Representative IF images of D21 CTRL, E:M and CTRL organoids exposed to conditioned media (CM) from pFG-SPM (referred to as CM) stained for C-PEPTIDE (green), TBX18 (blue) and Hoechst (grey). Scale bar, 50µm. **c,** Scatter plot showing the % of C-PEPTIDE^+^ cells relative to the total number of cells per organoid across conditions. Individual dots represent the mean value of biological replicates. Values shown are mean ± S.E.M. N= 3. (CTRL, n= 39; E:M, n= 66; CM, n=33). p=0.052 (paired t-test). **d,** Representative IF images of D21 CTRL, E:M and CM organoids stained for PDGFRB (green), COLLAGEN IV (COLIV) (red), PDX1 (blue) and Hoechst (grey). Scale bar, 50µm. **e,** Scatter plots showing the % of PDGFRB^+^ cells relative to the total number of cells per organoid across conditions. Individual dots represent the mean value of biological replicates. Values shown are mean ± S.E.M. N= 3. (CTRL, n= 11; E:M, n= 31; CM, n=28). **f,** RT-qPCR analysis of selected endocrine genes in CTRL and DOME organoids at D21. Data are shown as LOG2 ratio relative to undifferentiated iPSCs. Values shown are mean ± S.E.M. N=4.

To further assess the contribution of direct cell-cell contact to differentiation, we performed an additional experiment using an indirect co-culture set-up. Specifically, PP organoids were cultured in suspension in the presence of pFG-SPM encapsulated in Geltrex, as for dome cultures, without direct physical contact between the two cell populations throughout the duration of differentiation protocol (Fig. 4a). Under these conditions, cells were exposed to the same medium as in the direct E:M co-culture, but PP and mesenchyme remained physically separated. Such indirect co-culture was not sufficient to enhance the expression of endocrine transcription factors, such as *NKX6-1*, or *INS* expression at D21 (Fig. 4f). Together these results indicate that soluble factors alone do not recapitulate the effects of the direct E:M co-culture.

### iPSC-derived pancreatic mesenchyme recapitulates *in vivo* microenvironmental niches

Next, we focused on the mesodermal cellular fraction to define the intercellular communication networks and molecular programs that influence endocrine lineage commitment within the engineered E:M pancreatic organoids. Reclustering of the mesenchyme compartment from CTRL and E:M organoids (D11, D14 and D21) identified four distinct mesenchyme clusters and a small EC population (Fig. 5a). Organoid mesenchyme (O_Mes) cluster 1, characterized by *PODXL* and *CLDN7* expression, resembled mesothelial cells based on the mapping to *in vivo* foetal pancreatic tissue single-cell and spatial transcriptomics datasets (Fig. 5b-d). O_Mes 2 and 3 populations had less well-defined *in vivo* counterparts, broadly mapping to the *in vivo* M-SubMeso and proliferative cell clusters, respectively (Fig. 5b-d). O_Mes 4 cluster was characterized by the expression of pericyte markers, including *PDGFRB, RGS5, ITGA8,* together with a set of pancreatic mesenchyme-associated transcription factors (*e.g., FOXF1*, *NKX3-2*, *NKX2-5*) and signalling molecules, previously associated to pancreas development *(e.g., WNT5A, SLIT2)*^12^ (Fig. 5b; Supplementary Table S6). This population emerged primarily in E:M organoids and showed close transcriptional similarity to pericytes and M-ECM populations when benchmarked against *in vivo* human foetal pancreatic tissue (Fig. 5c-e and Extended Data Fig. 9a). Importantly, IF staining for ACTA2, PDGFRB, ITGA8, CSPG4 confirmed the presence of a comparable mesenchymal population in E:M organoids and *in vivo* in human foetal pancreatic tissue. These cells expressed the same vascular-associated marker set and occupied a peri-islet niche adjacent to INSULIN^+^ cells *in vivo* (Fig. 5f).

**Fig. 5.**
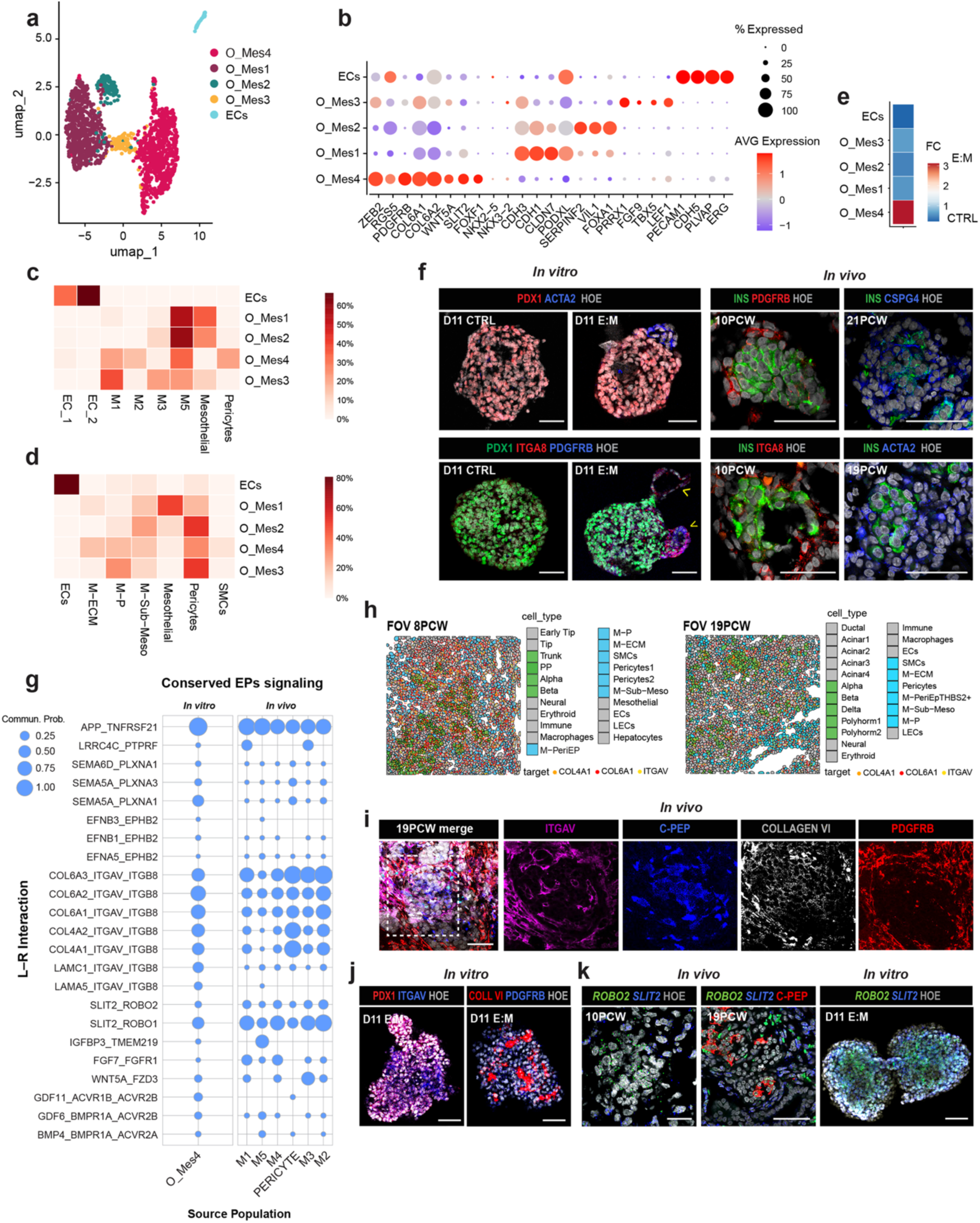
Remodelling of the mesodermal lineages in the E:M organoids. **a,** UMAP visualization of the mesoderm compartment from sc-RNASeq analysis on CTRL and E:M organoids. **b**, Dot Plot showing the expression of selected markers across populations identified. Dot size indicates the % of expressing cells; colour represents scaled AVG expression level (Supplementary Table S6). Abbreviations: ECs, Endothelial cells. O_Mes, Organoid Mesenchyme. **c**, Co-occurrence heatmap showing the correspondence between mesoderm *in vitro* labels and transferred labels from *in vivo* dataset^6^. **d**, Co-occurrence heatmap showing the correspondence between manually annotated labels and transferred labels from *in vivo* spatial mesoderm compartment from the CosMx dataset. **e**, Heatmap showing cell type fold change (FC) between co-culture (E:M) and control (CTRL) conditions. **f**, Left panels, representative IF images of D11 CTRL and E:M *in vitro* organoids stained for PDX1 (red), ACTA2 (blue), and Hoechst (grey) (Top); PDX1 (green), ITGA8 (red), PDGFRB (blue) and Hoechst (grey) (Bottom). Arrowheads indicate ITGA8^+^/PDGFRB^+^ cells at the periphery of E:M organoids. Right panels, representative IF images of *in vivo* human foetal pancreatic tissue stained for INSULIN (INS) and indicated mesenchymal/pericytes (ITGA8; PDGFRB; CSPG4) markers, and Hoechst (grey). Scale bars, 50μm. **g,** Conserved ligand-receptor (L:R) interactors that contribute to the endocrine progenitors (EPs) signalling from indicated cellular sources *in vitro* in D11 E:M organoids and *in vivo* pancreatic tissue. Analysis was performed using CellChat based on sc-RNASeq data from our organoids and publicly available datasets^6^. Dot size represents the calculated communication probability. **h**, Representative fields of view (FOV) of 8 PCW (left) and 19PCW (right) sections highlighting indicated cell populations and selected gene targets. **i**, Representative multiplex IF images on 19PCW human foetal pancreatic tissue stained for C-PEPTIDE (C-PEP) (blue), ITGAV (magenta), COLLAGEN VI (white), PDGFRB (red) and Hoechst (grey). Scale bars, 50μm. **j**, Representative IF images of D11 E:M organoids stained for PDX1 (red), ITGAV (blue), and Hoechst (grey) (Left) or COLVI (red), PDGFRB (blue), and Hoechst (grey) (Right). Scale bar, 50μm. **k,** Representative images of RNAscope multiplex fluorescent *in situ* hybridisation coupled to IF on human foetal pancreatic tissue or D11 E:M organoids for *ROBO2* (green) and *SLIT2* (blue) RNAs, C-PEPTIDE (red) and Hoechst (grey), at indicated foetal stages. Scale bars, 50μm.

To investigate how the O_Mes 4 mesenchymal population may influence endocrine differentiation, we examined the signalling pathways and adhesion molecules potentially mediating their interactions using CellChat. The analysis of cell-cell communication networks predicted O_Mes 4 as an active ‘sender’, exhibiting numerous predicted outgoing ligand–receptor (L:R) interactions with EPs, which were also conserved *in vivo* in foetal pancreatic tissue datasets^6^ (Fig. 5g). Among the inferred L:R interactions, some have previously been implicated in endocrine differentiation and islet morphogenesis in mouse, including Collagen IV and VI–Integrin^12,33–35^ interactions, BMP4^36^, EPH/Ephrin bidirectional signalling^37^ and SLIT-ROBO^37,38^, even though their roles in human endocrine development remain poorly understood. Other L:R interactions, such as LAMC-ITGAV/ITGB8, LNFN5-PTPRF, APP-TNFRSF21, TNM4-ADGRL1, have not yet been explored in this context.

Our spatial transcriptomics data enabled us to map the *in vivo* distribution of transcripts corresponding to the identified L:R pairs within the tissue, highlighting the spatial juxtaposition of mesenchymal populations, such as M-ECM and pericytes, with endocrine cells (Fig. 5h). Consistent with these findings, multiplexed IF analysis combined with RNAScope validated the spatial distribution of a sub-set of these L:R pairs in human foetal pancreatic tissue *in vivo* and in E:M pancreatic organoids *in vitro* (Fig. 5i,j). For example, at 19PCW COLLAGEN VI was present in PDGRB^+^ cells, while its receptor ITGAV in adjacent C-PEPTIDE^+^ beta-cells, consistent with a potential signalling interaction (Fig. 5i). A comparable spatial distribution pattern for both L:R components was observed in E:M organoids (Fig. 5j). Additionally, we validated the expression of *SLIT2* and its receptor *ROBO2* in C-PEPTIDE^+^ beta-cells in both human foetal tissue and in E:M organoids (Fig. 5k).

Together, these findings indicate that the iPSC-derived pFG-SPM acquires tissue-specific features through interaction with the pancreatic endoderm. Further supporting this conclusion, exposure of the starting pFG-SPM to the same defined set of pancreatic differentiation cytokines used in the organoid culture did not induce major changes in the expression of mesenchymal genes, as assessed by RT-qPCR (Extended Data Fig. 9b,c). The programmed O-Mes 4 resembles pancreatic mesenchyme populations found *in vivo* and exhibits a set of signalling modules, including guidance and ECM cues, that may contribute to the establishment of a supportive endocrine niche and promote beta-cell differentiation.

Next, we combined L:R inference with spatial analyses of the human foetal pancreas to further map mesenchymal signalling events within the microenvironment. CellChat L:R analysis identified a same set of guidance cues associated with neural repulsion, including SLIT2-ROBO, across pancreatic mesenchymal populations, predicting interactions not only with endocrine cells (Fig. 5g) but also with neural cells, both *in vivo* and *in vitro* (Extended Data Fig. 10a). Additionally, our CosMx niche analysis showed that trunk and endocrine progenitors preferentially co-localized with mesoderm-derived populations, particularly vascular-associated cells, whereas neural cells were largely excluded from these niches at both developmental stages analysed (Fig. 1d,g). Consistent with the niche organization, neural cells were located farther from endocrine cells than from vascular-associated populations, including pericytes (Extended Data Fig. 10b-d). In line with this *in vivo* spatial organization, we observed a pronounced reduction in off-target neural-like cells *in vitro* in the E:M organoids upon addition of pFG-SPM (Fig. 2e) along with a reduced expression of neural marker genes (Extended Data Fig. 10e). Similarly, the addition of SLIT2 ligand to CTRL organoid cultures phenocopied this effect, resulting in reduced expression of neural-specific markers (Extended Data Fig. 10f). Together, these findings suggest that pancreatic mesenchymal populations contribute to endocrine niche organization, while limiting neural–endocrine proximity.

## Discussion

Here, we combined spatial transcriptomics of the developing human pancreas with an iPSC-derived multilineage organoid system to explore cellular and molecular features of the human endocrine niche. Our spatial mapping of the developing human pancreas at single-cell resolution reveals a more complex mesenchymal landscape than previously recognized. Whilst we confirm the presence of vascular, proliferative, and mesothelial populations identified in lower-resolution studies^11,39^, our analysis includes an earlier developmental stage (8PCW), not previously available, and reveals additional mesenchymal populations with different features and occupying spatially distinct tissue niches. Tissue niches are dynamic and exhibit distinct cellular compositions, with different mesenchymal populations associated with specific pancreatic cell types, suggesting that they may provide specialised cellular environments. For example, the M-ECM population was enriched in the endocrine niche, while the M-CCL3 was more abundant in the duct/acinar niche. Conversely, some mesenchymal populations were shared across multiple niches, although whether they adopt distinct gene expression profiles depending on their niche remains to be addressed. Further studies will also be required to determine the functional roles of the identified niches and to elucidate the underlying ligand–receptor interactions between distinct cell populations.

The strong concordance between our spatially-defined CosMx populations and sc-RNASeq datasets^6,7,20^ validates our cell atlas and provides a valuable framework for investigating cell-cell interactions in the human foetal pancreas. The integration of high-quality data from other datasets will enable the extension of our atlas beyond development to adult stages and disease states, such as diabetes. Therefore, functional studies of these cellular interactions will not only provide mechanistic insight into human pancreatic development but may also unveil niche alterations associated with disease. For example, disruption in niche composition and cellular localisation have been reported in cystic fibrosis, highlighting the potential contribution of altered cellular microenvironments to disease pathogenesis^40^.

Faithfully recapitulating cellular heterogeneity and tissue architecture of human organs is a key limitation of organoid technology. Our spatial atlas and sc-RNASeq datasets have provided a benchmark for establishing a multilineage pancreatic organoid model that combines iPSC-derived PPs with pFG-SPM. This model recapitulates key features of the endogenous composition of the developing human pancreas, especially the endocrine lineages. Similarly, the pFG-SPM incorporated to the organoid culture undergoes remodelling, giving rise to a heterogeneous mesenchyme with pancreatic tissue resident features that closely resemble some *in vivo* populations. We found that off-target spontaneous mesenchyme represents only a small fraction of the culture under control conditions (< 5%) and does not overlap with identifiable *in vivo* counterparts. Notably, our multi-lineage pancreatic organoid model underscores a bidirectional exchange between endoderm and mesoderm. The interaction of programmed pFG-SPM with pancreatic endoderm is required for mesodermal-niche remodelling and acquisition of tissue-specific signatures. At the same time, mesodermal niche cues direct regionally appropriate pancreatic differentiation, enhancing lineage fidelity and improving beta-cell differentiation.

We map a dynamic endocrine niche in which the pancreatic mesenchyme emerges as multifunctional orchestrators, coordinating signalling and ECM interactions with endocrine cells, while also providing repulsive guidance cues, such as SLIT, Semaphorin, and Ephrin signalling, that target both endocrine and neural cells. Collectively, our findings support a model in which a coordinated program of repulsive signals might shape the spatial organization of the developing endocrine niche and islet–neural interactions, defining physical boundaries of emerging islets while delaying innervation. Consistent with this model, a temporal relationship, whereby neural recruitment follows the establishment of the vascular niche, has been described in the mouse pancreas^41,42^ and is supported by recent observations in the human postnatal pancreas^43^.

Together, our spatial atlas and multilineage pancreatic organoid system establish an advanced human platform that faithfully recapitulates key features of the foetal endocrine niche, enabling mechanistic studies of human pancreatic development, metabolic homeostasis, and disease, such as diabetes, and, ultimately, opening new avenues for the engineering of transplantable tissues for diabetes cell-based therapies.

## Acknowledgement

We thank all the members of the Spagnoli laboratory for their useful comments and suggestions on the study. We thank Heiko Lickert (Helmholtz, Munich) for the hiPSC line HMGUi001-A2. We thank Rosamond Nuamah, Jelmar Quist, Roman Laddach, Ciro Chiappini and Anita Grigoriadis at The Spatial Biology Facility (Innovation Hub, Guy’s Cancer Centre, King’s College London) for their assistance in performing the CosMx^TM^ SMI single-cell Spatial Transcriptomics. We thank Dasha Freydina at the Single Cell Omics Platform (CIBCI, King’s College London) for performing the library preparation of the GEM-X Flex sc-RNASeq and her assistance throughout the process and the Advanced Cytometry Platform (Flow Core) at King’s College London. We are grateful to the Human Developmental Biology Resource (http://hdbr.org) for the help in collecting the pancreatic foetal tissue.

## Author contributions

Conceptualization: F.M.S., G.G., A.T.C. Data curation: G.G., A.T.C., F.M.S. Funding acquisition: F.M.S. Investigation: G.G., A.T.C., H.W, R.M.G. Methodology: G.G., A.T.C., H.W., R.M.G., S.M., L.H.A., M.P., M.S, F.F., A.V. Project administration: F.M.S. Supervision: F.M.S. Writing—original draft: G.G. Writing—review and editing: F.M.S, G.G and A.T.C.

## Online Methods

### Cell Lines and Cell Culture

Human iPSC cell line HMGUi001-A2 (sex: female) was kindly provided by Heiko Lickert (Helmholtz, Munich) and authenticated by karyotyping. Human iPSC cell line AICS-0090 from the Allen consortium (sex: male) was obtained from the Coriell Institute for Medical Research. Human iPSCs were maintained on Geltrex-coated (Invitrogen) plates in E8 media. The medium was changed daily, and cells were passaged every 3-4 days as cell clumps or single cells using EDTA (Invitrogen) or Accutase (Invitrogen), respectively. The medium was supplemented with Rho-associated protein kinase (ROCK) inhibitor Y-27632 (Sigma) when iPSCs were thawed or passaged as single cells.

### Directed differentiation of human iPSCs into pancreatic cells

Human iPSC pancreatic differentiation was carried out following the protocol of Russ *et al*.^27^. Briefly, iPSCs were dissociated using Accutase and seeded at a density of 5.5×10^6^ million cells per well in a 6 well-ultralow attachment plates (CellStar) in E8 medium supplemented with ROCK inhibitor (10μM), Activin A (10ng/ml), Heregulin-b1 (10ng/ml) and Glutamax and placed onto an orbital shaker rotating at 100rpm for suspension culture. Cells remained in 3D suspension on the orbital shaker for the duration of the protocol. From day (D) 1 to D5 cells were in RPMI media and from D6 to D21 in DMEM with 25mM Glucose. Media was changed daily as previously described by Russ *et al*.^27^. See Supplementary Table S7 for cytokines and differentiation protocol. For pathway perturbation experiments, recombinant SLIT2 (100 ng/mL) was added to the culture medium on D9, D10, and D11 of CTRL organoids.

### Directed differentiation of human iPSCs into posterior foregut splanchnic mesoderm (pFG-SPM)

Directed differentiation of iPSCs into pFG-SPM was carried out as previously reported in Kishimoto *et al*^22^. Briefly, iPSCs were dissociated using Accutase, seeded at a density of 4×10^5^ onto Geltrex-coated 6-well plates (Corning) and differentiated as a 2D monolayer. First, primitive streak was induced by culturing cells in a basal media composed of DMEM/F12, 15mM HEPES, supplemented with Activin A, CHIR, BMP4, bFGF, and PIK90 for 24 hours. Next, for the induction of the lateral plate mesoderm cells were then exposed to A8301, BMP4, retinoic acid and C59 for 24 hours and for pFG-SPM generation, cells were cultured in A8301, BMP4, FGF2, and RA from day 2 to day 4. See Supplementary Table S7 for cytokines and differentiation protocol.

To fluorescently label pFG-SPM cells, lentiviral particles were prepared from the U6-sgRNA/EF1a-mCherry lentivirus vector (Addgene, plasmid no. 114199). Cells were transduced with lentiviral particles at a MOI of 10 on day 3 of differentiation, 24 h before establishing the co-culture. After 24 h, cells were detached, counted, and added to the PP cell culture. Successful mCherry labelling was confirmed by fluorescence microscopy in both 2D cultures and 3D co-cultures 24 h after transduction.

### Co-culture of pancreatic progenitors with pFG-SPM to generate multilineage organoids

Pancreatic Progenitors (PP) cells were combined with D4 pFG-SPM. Briefly, pFG-SPM cells were dissociated into single cells. Simultaneously, PP clusters from one well of 6-well plate were collected using a 0.2% BSA/ PBS coated 10ml pipette and divided into two Falcon tubes. 1.75×10^6^ pFG-SPM cells were added to each Falcon tube containing the PP clusters. Falcon tubes were centrifuged for 3 minutes at 300g. Mixed cell populations were gathered using a 10ml 0.2% BSA/PBS coated pipette, pipetted gently up and down to mix and then dispensed back into one well of a 6-well ultralow attachment plate on the orbital shaker. Medium composition for multilineage organoids is a 50:50 mix of pancreatic differentiation medium of appropriate stage of differentiation with the pFG-SPM medium (DMEM/F12 supplemented with B27, N2, and Glutamine, without the addition of BMP4, A83-01, FGF2, RA, Wnt-C59 cytokines) (Supplementary Table S7).^22^ The basic condition culture medium (BM) consists of DMEM/F12 supplemented with B27, N2, and Glutamine.

For indirect co-culture condition with pFG-SPM conditioned media (CM), culture medium from pFG-SPM cells at D4 was collected, centrifuged at 300g for 5 minutes, passed through a 0.22μm filter and stored at −80°C. Thawed CM was then mixed 1:1 with fresh pancreatic differentiation medium, corresponding to the appropriate stage of differentiation, and applied to cells from D9 (PP stage) until D21.

For indirect co-culture condition with Geltrex encapsulation, six 50-µL domes were generated per well of a six-well ultra-low-attachment plate. A total of 1.75 × 10^6^ pFG-SPM cells were resuspended in 300 µL Geltrex and distributed equally across six domes positioned around the well. PP cells were maintained in the centre of the well.

### Library preparation and sc-RNA sequencing

Organoids were collected with a pipette coated with 0.5% BSA-PBS, washed and centrifuged for 1 minute at 300g, before being transferred into 1.5ml Eppendorf tubes. 600ul of Accutase was added and Eppendorf tubes were incubated in a thermomixer set to 23°C at 600rpm for 15 minutes. Following this, 600ul of DMEM/F12 was added to each tube and cells were then centrifuged for 1 minute at 300g, supernatant was then removed and the cell pellet was resuspended in DMEM F21 media and passed through a 70um cell strainer. Simultaneously, cell viability was assessed using AO/PI staining and counted using the Countess 3 FL automated cell counter (ThermoFisher). Fixation was performed using the ‘Fixation of Cells & Nuclei’ for GEM-X Flex Gene Expression kit (10x Genomics; CG000785, Rev A), a formaldehyde-based fixation approach. Briefly, samples were fixed for 22 hours overnight (ON) at 4°C. Cells were then quenched using chilled Quenching buffer and counted again with AO/PI using the Countess automated cell counter. Long-term storage procedure at −80°C was then followed – adding the enhancer (10x Genomics PN-2000482) and 50% glycerol to the quenching buffer solution. Samples were then transferred to the Single Cell Omics Platform at KCL for library preparation and multiplexing using the GEM-X Flex GEM & Library Kit (PN-1000782) with the GEM-X Flex Human Transcriptome Probe Kit (PN-1000785). 8 samples were multiplexed and combined into two pools (representing 4 multiplexed samples each) on separate 25B lanes. Novogene performed the GEM-X Flex 10x probe-based sequencing; 20,000 cells per sample, 30-35,000 reads per cell (Premade Library Sequencing 840Gb ∼2,800 million reads per library, Illumina Novaseq X Plus | PE150). GEM-X Flex technology was chosen as it allowed us to fix our samples at each timepoint and then perform the library prep and sequencing together to avoid possible batch effects.

### Processing of sc-RNASeq data

Raw multiplexed sequencing reads were demultiplexed and aligned to GRCH38 using the nf-core/sc-RNASeq v2.7.0 and the Cellrangermulti (Cellranger:9.01) pipeline with default parameters. Barcode references used for demultiplexing were: BC001, BC002, BC003, BC004. Samples for each barcode were: BC001 (D4 pFG-SPM, D14 Control), BC002 (D9 PP, D14 E:M Co-culture), BC003 (D11 Control, D21 Control), BC004 (D11 E:M Co-culture, D21 E:M Co-culture). Downstream analysis was completed with Seurat (v.5.3.0).

For quality control, low-quality cells with a number of features lower than 1000, a number of gene counts below 1000 and a mitochondrial ratio higher than 15% were removed. Doublets were identified with the package DoubletFinder(v2.0)^44^ and subsequently removed. Count matrix was then normalized and scaled using the SCTransform() Seurat function^45^. After performing Principal Component Analysis (PCA) for dimension reduction, batch effects between different samples was corrected using Harmony^46^. Harmony dimensions were then used to generate a low-resolution clustering using FindNeighbours() and FindClusters() Seurat’s functions. After that, each low-resolution cluster was subset from the full object, and subclustered by calculating new PCA and Harmony dimensions. Similarly, Endocrine cells from the Endoderm compartment (Endo, Endo Immature and EP) were subset and analysed separately. To visualise the data, we used uniform manifold approximation and projection (UMAP). Differentially expressed genes were calculated with the FindAllMarkers() Seurat’s function.

### sc-RNASeq *in vitro* benchmark to *in vivo* sc-RNASeq and spatial datasets

To benchmark *in vitro* clusters to their *in vivo* counterparts, we used the publicly available dataset Ma et al.^6^ (OMIX001616-01). From this dataset, we used cells belonging to 7 to 11PCW stages and processed and annotated them following the same pipeline as written in the previous section. For each major organoid compartment, we asked which *in vivo* clusters resembled the annotated *in vitro* clusters. For this, related clusters from the *in vivo* data were subset (epithelial cells for the endoderm compartment, mesenchymal and mesothelial cells for the mesoderm compartment and endocrine cells for the endocrine subset) and labels were transfer using the TransferAnchors() and TransferData() Seurat’s functions. Label co-occurrence was visualized using a heatmap. For the endocrine and mesoderm compartments we also compared the *in vitro* clusters to the endocrine and mesenchymal clusters annotated in the 19PCW and 8PCW, respectively, CosMx datasets (see below).

### Cell–cell communication analysis

Cell–cell communication networks were inferred using CellChat (v2.2.0)^47^ to identify ligand–receptor interactions present both in the *in vitro* co-culture system at Day 11 and a published *in vivo* human pancreatic dataset (Ma et al., 2024). CellChat objects were built independently for each dataset using normalized gene expression matrices and cell-type annotations, and ligand–receptor interactions were inferred using the CellChatDB database, with communication probabilities calculated from ligand and receptor expression across interacting populations. Interactions between endocrine progenitors (EPs) and mesenchymal populations were extracted from both *in vitro* and *in vivo* datasets. Ligand–receptor pairs detected in both datasets were classified as conserved, while those detected exclusively in one dataset were classified as *in vivo*-specific or *in vitro*-specific accordingly.

### Human Tissue

Human embryonic and foetal pancreas were provided by the Joint MRC/Wellcome Trust (grant#MR/X008304/1 and 226202/Z/22/Z) Human Developmental Biology Resource (http://hdbr.org) with appropriate maternal written consent and approval from the Newcastle and North Tyneside NHS Health Authority Joint Ethics Committee (23/NE/0135) and London Fulham Research Ethics Committee (23/LO/0312). The HDBR is regulated by the UK Human Tissue Authority (HTA; www.hta.gov.uk) and operates in accordance with the relevant HTA Codes of Practice. Human pancreatic tissue samples (gender not established) were fixed overnight in 4% PFA and then processed for sectioning, CosMx^TM^ spatial molecular imaging (SMI), immunostaining and imaging at King’s College London. All work was undertaken in approval of the HDBR Steering Committee to the Spagnoli lab. at King’s College London, UK (License #200523).

### CosMx^TM^ spatial molecular imaging (SMI)

CosMx^TM^ technology [CosMx^TM^ Human Universal Cell Characterization RNA Panel (1000-plex); Nanostring] was applied to 3 FFPE samples (8PCW, 15PCW, 19PCW); four sections per sample were analysed within an area of 20mm x 15mm. The following cell surface markers were used for morphology visualization and segmentation: PanCK, CD45, CD298/B2M antibodies and DAPI, as nuclear counterstaining. FFPE tissue sections were prepared for CosMx^TM^ SMI profiling as previously described by Shanshan *et al.*^48^. Briefly, FFPE blocks were sectioned at 5um thickness on Superfrost^TM^Plus Adhesion Microscope Slides (Epredia; 021624-9), dried at 37°C overnight at an angle no greater than 45 degrees. Slides were then stored at 4°C with a desiccant bag before being handed over to the Spatial Biology Facility at KCL for further processing and loading into the instrument. FOV selection took place whereby the entirety of every tissue section of each sample was selected.

### CosMx™ SMI sample segmentation, quality control, downstream processing

Segmentation of the image was carried out on the AtoMx platform using the CellPose algorithm Panel_A. After segmentation, z-stack cleaning was performed using the NoButter pipeline, as previously described by O’Hora *et al.*^49^. Briefly, the transcripts located in the top zstack of each section (selected due to not being associated with the tissue section / more likely to be noise and give false readouts), were removed. After which the new GEX matrix was created. Flat files were then exported from the AtoMx platform into R Studio whereby the unique cell ID and FOV ID was combined within the metadata. This allowed visualization of FOV position, tissue section position within the slide, cell position and mean expression level of B2M, PanCK, CD45, and DAPI. Sections from each stage were analysed separately. First, low-quality cells expressing less than 3 genes and 5% of negative probes were removed. Normalization was carried out using the function SCTransform(). After that we performed PCA and performed a low-resolution clustering. Each cluster was then subset and subclustered independently and annotations were imported into the full object.

To map dorsal and ventral pancreas regions of the 8PCW samples, areas were manually drawn over the CosMx sections using the gatepoints package (v.0.1.5). Cellular composition differences between both regions were calculated using each section as a replicate and the edgeR package (v.4.6.3)^50^. We performed two comparisons: one Dorsal *vs* Ventral cell abundances within the epithelial compartment and one comparing differences among non-epithelial cells.

### Niche Analysis

To identify tissue domains at 8PCW and 19PCW we used BANKSY^21^. Spatial weights were calculated using 15 neighbours, the Azimuthal Gabor filter, and gaussian decay. We used lambda value of 0.9, prioritizing the expression profile of its neighbours over its own. To identify niches, we then run PCA and Leiden clustering with a resolution of 0.7. At 19PCW one niche contained cells at the corners of FOV, was considered a technical artifact and was therefore removed.

### CosMx and sc-RNASeq data integration

To integrate sc-RNASeq and CosMx datasets, we matched 8PCW sections with 8PCW cells from Ma et al.^6^ sc-RNASeq dataset, and 19PCW sections with 18PCW cells from the Migliorini et al.^7^ dataset. In the case of the Migliorini dataset, we used the annotation provided by the original study, whilst in the case of the Ma dataset, cells were reclustered and reannotated to identify mesenchymal subpopulations. To integrate both modalities, first we removed features that were not present in both datasets. After SCT normalisation, spatial and sc-RNASeq datasets were integrated using Harmony. We then used the Harmony calculated space to transfer sc-RNASeq labels into the CosMx cells. Specifically, we used Python’s scikit-learn (v.1.5.2) implementation of K-means to obtain the 5 closest sc-RNASeq neighbours for each CosMX cell and transferred the most frequent label. The same procedure was followed to transfer the average gene expression from the 3000 more variable features of the 5 closest sc-RNASeq neighbours into CosMx cells.

### Immunofluorescence analysis on cells, organoids and tissue sections

iPSC-organoids were collected into Eppendorf tubes, fixed with 4% PFA for 20 minutes, washed with 0.5% BSA/PBS solution, and equilibrated within a 15% sucrose solution for 1 hour. Organoids were then encapsulated in a 2.5% low melting point agarose (Sigma-Aldrich) 15% sucrose PBS solution and transferred into embedding moulds filled with OCT compound (Tissue-Tek, Sakura). OCT blocks were then sectioned at 12 μm thickness. For Formalin-Fixed Paraffin-Embedded (FFPE) tissue, slides were incubated at 56° C for 1-2 hours, before xylene deparaffinization and rehydration in PBS.

For immunofluorescence (IF) staining, slides were placed in an oven at 37°c for 1 hour and permeabilised with 0.5% Triton X-100 PBS solution before antigen retrieval by boiling slides for 20 minutes in Citrate buffer (Dako). Slides were then permeabilised with 0.5% Triton X-100 PBS, before being incubated in TSA (Perkin Elmer) blocking buffer for 1 hour at room temperature (RT). Incubation in primary antibody solutions (3% Donkey Serum, 3% BSA, 0.1% Triton X-100) at the appropriate dilution (Supplementary Table S7) was carried out overnight (ON) at 4°C. Slides were then washed with 0.5% Triton X-100 PBS solution 3 times before being incubated in secondary antibody solutions (3% Donkey Serum, 3% BSA, 0.1% Triton X-100) (Supplementary Table S7) with Hoechst 33342 (Invitrogen, #H1399) at a concentration of 250ng/ml for 1 hour at RT. Slides were then washed with PBS solutions and mounted with Dako Fluorescent Mounting media (Dako, #S3023). Image acquisition was performed using Zeiss Discovery and Zeiss LSM 900 laser scanning confocal microscopes.

For IF on pFG-SPMs cultured as 2D monolayers, cells were washed with PBS and fixed in 4% PFA at RT for 20 minutes. Samples were then incubated with 3% Donkey serum/0.1% Triton X-100 PBS blocking buffer for 30 minutes at RT. Primary antibodies were diluted within fresh blocking solution at the appropriate concentration (Supplementary Table S7) and samples were incubated ON at 4°C. Cells were then washed 3 times with 0.1% Triton PBS before being incubated in the secondary antibody solution (3% Donkey serum, 0.1% Triton X-100 PBS plus 250ng/ml Hoechst) (Supplementary Table S7), for 30 minutes at RT. Samples were washed again with 0.1% Triton X-100 PBS, before being placed in PBS for imaging. Image acquisition was performed with an Operetta CLS (Perkin Elmer).

### Iterative indirect immunofluorescence imaging (4i) on tissue sections

Multiplexed immunofluorescence was performed on 20 WPC human pancreas tissue sections using the 4i (iterative indirect immunofluorescence) protocol, as described previously^51^. Briefly, OCT-embedded sections underwent antigen retrieval in Dako Target Retrieval Solution (20 min), permeabilization in PBS with 0.5% Triton X-100, and blocking in PBS with 4% BSA and maleimide (1 h). Sections were incubated with primary antibodies overnight at 4°C, followed by species-specific fluorescent secondary antibodies, DAPI nuclear counterstaining, and mounting in 4i imaging buffer.

For iterative staining, antibodies were eluted between rounds by immersing slides in distilled water to remove coverslips, followed by three sequential 10-min incubations in elution buffer with an interposed PBS wash; complete antibody removal was confirmed by the absence of residual fluorescence. Sections were then re-blocked for 1 h and re-stained as described above for each subsequent round. All images were acquired on a confocal microscope using identical acquisition settings across rounds.

Iterative 4i staining was performed across three rounds: Round 1 (ITGAV; Abcam ab97588, C-PEPTIDE; Cell Signalling 4593S), Round 2 (PDGFRB; Abcam ab32570), Round 3 (COL6A1; Abcam ab182744); representative images shown display ITGAV, C-PEPTIDE, PDGFRB, and COL6A1.

### RNAscope Multiplex Fluorescent *In Situ* Hybridization and Immunofluorescence

RNAscope Multiplex Fluorescent V2 *in situ* hybridization (Advanced Cell Diagnostics) was performed on frozen tissue sections and fixed organoids according to the manufacturer’s protocol, with an adapted protocol used for organoids as previously described^52^.

Frozen sections and fixed organoids were air-dried at −20°C for ≥1 h, treated with RNAscope Hydrogen Peroxide (10 min, RT), and subjected to heat-induced antigen retrieval in 1× Target Retrieval Reagent (5 min), followed by dehydration in 100% ethanol. Sections were then treated with Protease Plus (15 min, 40°C) prior to probe hybridization.

C1 probes were used alone or as carriers for C2 probes (1:50, C2:C1), applied for 2 h at 40°C. RNAscope probes used were: RNAscope™ Probe - Hs-SLIT2 (cat #466221) and RNAscope™ Probe - Hs-ROBO2-C2 (cat # 502341-C2). Signal was amplified sequentially using AMP1–AMP3 reagents, and channel-specific HRP (HRP-C1/C2) was developed and detected by tyramide signal amplification (TSA Plus or Opal fluorophores, 1:750, 30 min, 40°C), with fluorophore choice matched to target transcript abundance (fluorescein for high-abundance, Cy3/Cy5 for low-abundance targets). Between channels, HRP was inactivated using RNAscope HRP blocker before sequential detection of the next probe with a spectrally distinct fluorophore.

Following RNAscope detection, tissue sections were blocked in 3% BSA/PBS (15 min, RT) and incubated with primary antibody overnight at 4°C, followed by fluorescently labelled secondary antibody (1 h, RT) and mounting in Dako Mounting Medium.

### RT-qPCR analysis

RNA was isolated from organoids and iPSCs using the High-Pure RNA Isolation Kit (Roche; 11929665001). RNA was then transcribed to cDNA using the Transcriptor First Strand cDNA Synthesis Kit (Roche; 0489703001). Quantitative real-time PCR (qPCR) was performed using FastStart Essential DNA Green Master (Roche; 06924204001) with Syber-Green probes (Supplementary Table S7) within LightCycler 480 Multiwell plate 96 (Roche; 04729692001) on a LightCycler 96 machine (Roche).

### C-PEPTIDE ELISA assay

Supernatants were collected across all conditions at the D21 timepoint, filtered using a 0.22μm filter and stored at −80°C until use. C-PEPTIDE ELISA was conducted using the Mercodia Ultrasensitive C-PEPTIDE ELISA kit (10-1141-01), absorbance was read at 450nm filter using a GloMax Discover Microplate Reader (Promega). Concentrations (pmol/L) were extrapolated using the provided 5 calibrator concentrations. An XY data table with the calibrator concentrations and corresponding absorbance values was generated with our sample absorbances, we then selected “Interpolate a Standard Curve”, and GraphPad Prism calculated the unknown sample concentrations. GraphPad Prism version 10.0.0 (GraphPad Software).

### Image analysis, statistics and reproducibility

Images were processed for visualization purposes using Fiji^53^. Quantifications of % of positive cells were performed using QuPath-0.5.1-x64^54^. Briefly, the number of positive cells was calculated either by using the positive cell detection, using Hoechst-positive nuclei or cell detection was first used to detect Hoechst-positive nuclei and then object classification; “create single measurement classifier” was used to detect positive cells in the 488, 555, and 647 channels. “load object classifier” was then applied to the image and single, and double positive cells were calculated. Measurements were then exported into excel and the % was calculated using the number of total cells detected in the image. Data representation, statistical analyses, and plotting of regression lines were performed using GraphPad Prism (version 10.0.0). Unless stated otherwise, data are shown as mean ± standard deviation (SD) or standard error of mean (S.E.M.) and *N* numbers refer to biologically independent replicates. Statistical significance (p<0.05) was determined as indicated in figure legends using paired two-tailed Student t-test, or One-way Anova, for comparison between more than two samples. Statistical analysis was conducted on all collected samples and data. No statistical method was used to predetermine the sample size. No data were excluded from the analyses. The experiments were not randomized. Investigators were not blinded to allocation during experiments and outcome assessment.

## Data and Materials availability

All data needed to evaluate the conclusions in the paper are present in the paper and/or the Supplementary Information. sc-RNASeq and CosMx spatial transcriptomics data generated in this study have been deposited in the Gene Expression Omnibus (GEO) under accession number GSE313331. Public sc-RNASeq datasets were collected from GEO (GSE230403, GSE167880), OMIX database: OMIX001616-01 and ArrayExpress database: E-MTAB-10187.

