## Extended data file for "Mesodermal-niche interactions direct specification and differentiation of pancreatic endocrine cells in human multilineage organoids"

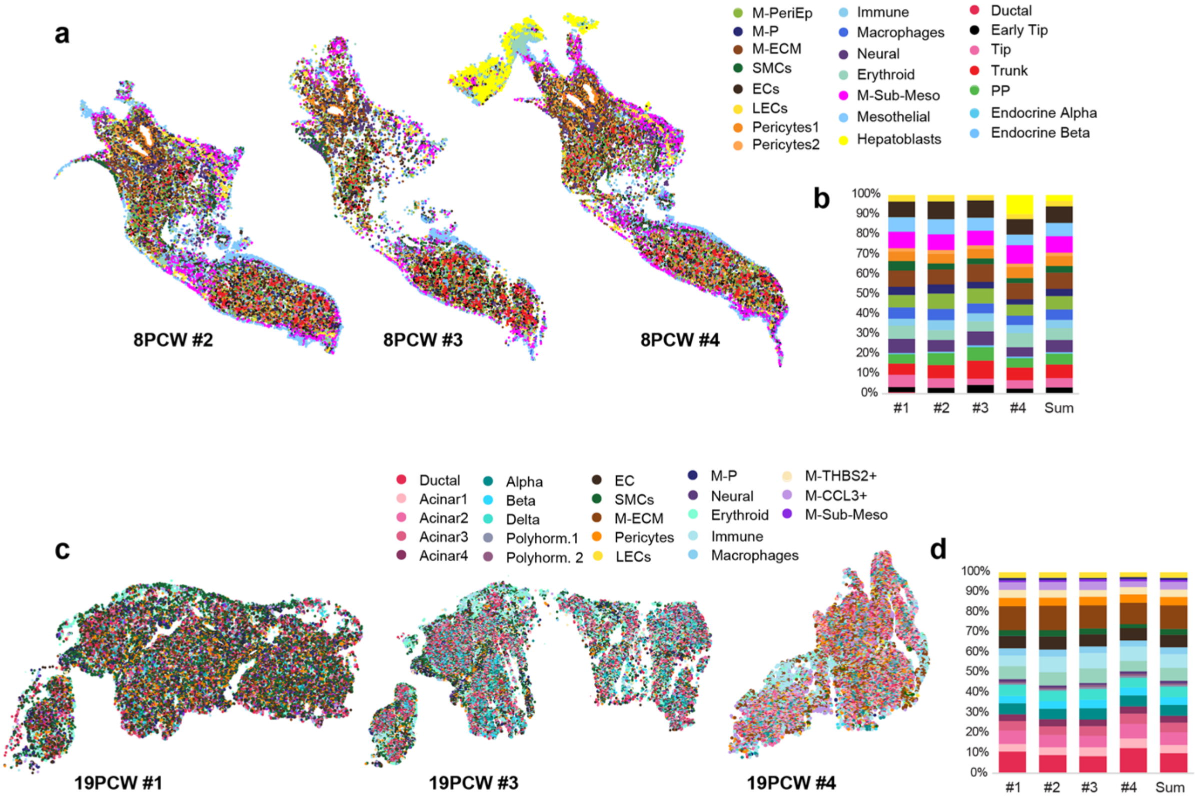


**Extended Data Fig. 1 | CosMx cell annotations are reproducibly mapped across tissue sections. a,** Spatial distribution of annotated cell populations from CosMx™ SMI data on 8PCW human pancreatic tissue sections. Each colour represents a cell type. **b**, Cell-type proportions in each of the 8PCW pancreatic tissue sections. **c,** Spatial distribution of annotated cell populations from CosMx™ SMI data on 19PCW human pancreatic tissue sections. Each colour represents a cell type. **d**, Cell-type proportions in each of the 19PCW pancreatic tissue sections.

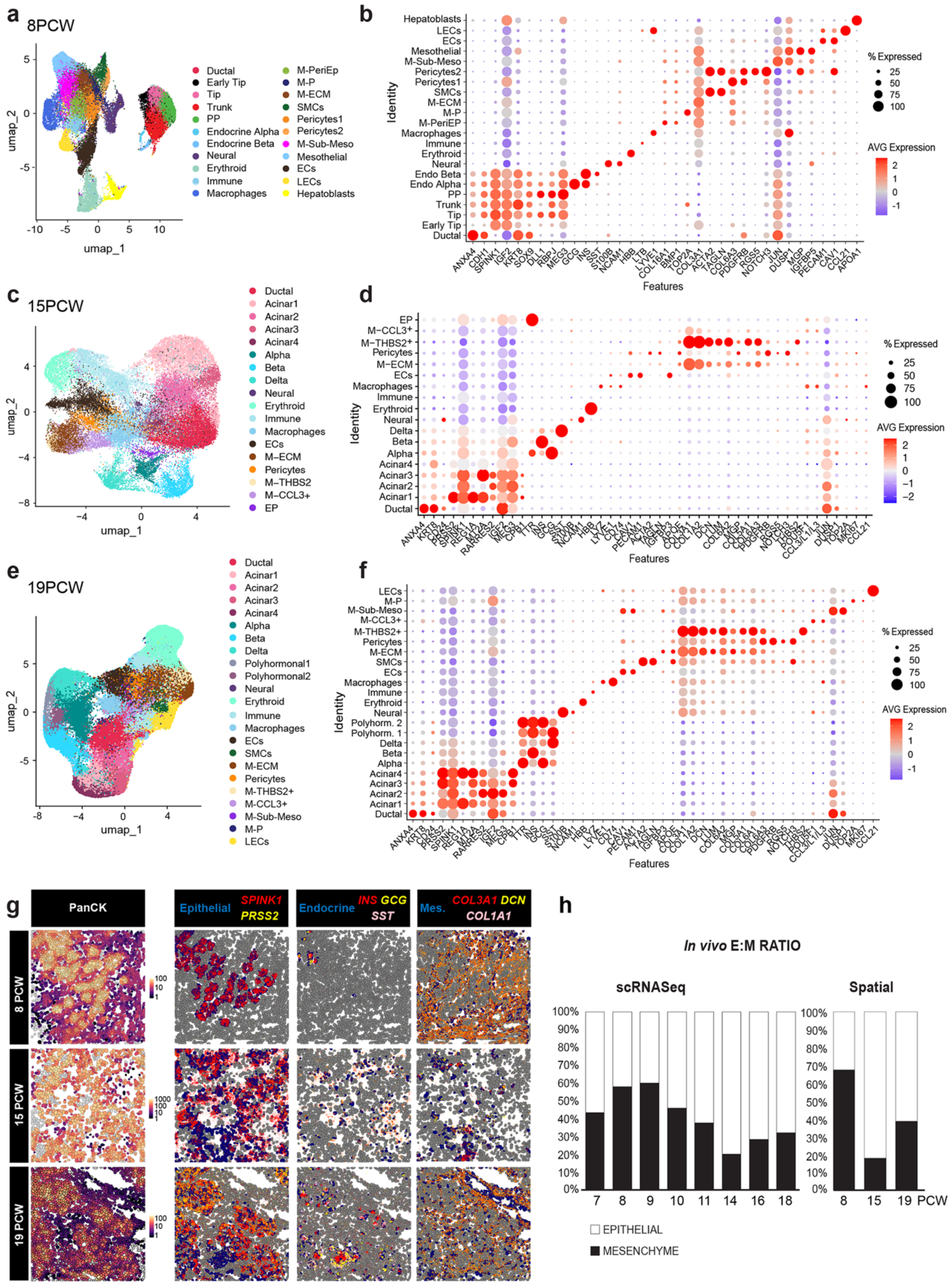

**Extended Data Fig. 2 | Cell populations of the human foetal pancreas at 8, 15 and 19 PCW. a,** UMAP visualization of 8 PCW pancreatic cells identified in the CosMx™ SMI dataset. **b**, Dot plot showing the expression of selected markers across cell populations identified at 8PCW. Dot size indicates the percentage (%) of expressing cells; colour represents scaled average (AVG) expression level (Supplementary Information Table S1). Abbreviations: PP, Pancreatic Progenitors; M-, Mesenchyme; PeriEp, Peri-Epithelial; P, Proliferative; SMCs, Smooth Muscle Cells; Sub-Meso, Sub-Mesothelial; EC, Endothelial Cells; LECs, Lymphatic Endothelial Cells. **c,** UMAP visualization of 15PCW pancreatic cells identified in the CosMx™ SMI dataset. **d,** Dot plot showing the expression of selected markers across cell populations identified at 15PCW. Dot size indicates the % of expressing cells; colour represents scaled AVG expression level (Supplementary Information Table S2). Abbreviations: EC, Endothelial Cells; M-, Mesenchyme; EP, Endocrine Progenitors. **e,** UMAP visualization of 19PCW pancreatic cells identified in the CosMx™ SMI dataset. **f**, Dot plot showing the expression of selected markers across cell populations identified at 19PCW. Dot size indicates the % of expressing cells; colour represents scaled AVG expression level (Supplementary Information Table S3). Abbreviations: ECs, Endothelial Cells; SMCs, Smooth Muscle Cells; M-, Mesenchyme; Sub-Meso, Sub-Mesothelial; P, Proliferative; LECs, Lymphatic Endothelial Cells. **g**, Representative fields of view (FOV) of 8, 15 and 19 PCW sections, showing selected gene markers and their expression in indicated cell populations. Left panel, Pan-Cytokeratin (PanCK) staining marking epithelial cells in the CosMx™ SMI platform. Scale bar indicates mean fluorescent intensity. **h**, Epithelial-mesenchymal cell proportions across different foetal stages measured from sc-RNASeq datasets Ma et al.[1] (7 to 11 PCW) and Migliorini et al.[2] (14 to 19PCW) and CosMx™ SMI spatial data.

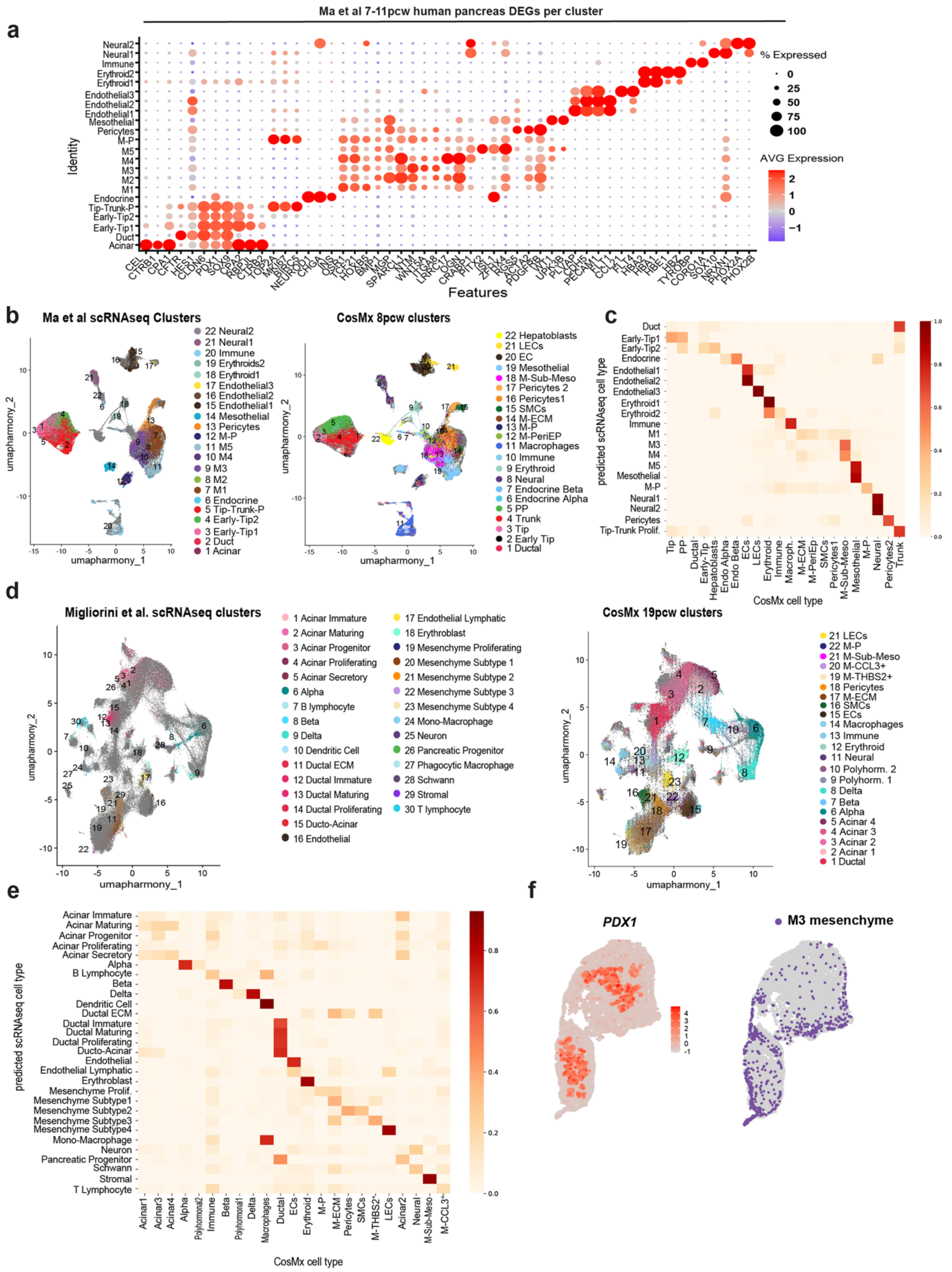


**Extended Data Fig. 3 | Systematic benchmarking of CosMx spatial data with sc-RNASeq datasets from human pancreatic tissue at equivalent developmental stages. a,** Dot plot showing the expression of selected marker genes across cell populations identified after reanalysis of published sc-RNASeq dataset of 7-11 PCW human foetal pancreas from Ma *et al.*[1]. Dot size indicates the % of expressing cells; colour represents scaled AVG expression level. **b,** UMAP plots showing clustering of integrated sc-RNASeq profiles of 7-11 PCW foetal pancreatic tissue obtained from Ma *et al.*[1] and spatial profiles from our CosMx 8PCW dataset. Left plot highlights cells from Ma *et al.*[1] sc-RNASeq; right plot, the CosMx cells. **c,** Co-occurrence heatmap showing predicted correspondences between 8PCW CosMx annotation and transferred labels from Ma *et al.*[1]. Colour scale represents the scaled overlap between datasets. **d,** UMAP plots showing clustering of sc-RNASeq profiles of 14-18 PCW foetal pancreatic tissue obtained from Migliorini *et al.*[2] and spatial profiles from our CosMx 19PCW dataset. Left plot highlights cells from Migliorini *et al.*[2] sc-RNASeq; right plot CosMx cells. **e,** Co-occurrence heatmap showing predicted correspondences between 19PCW CosMx annotation and transferred labels from Migliorini *et al.*[2] Colour scale represents the scaled overlap between datasets. **f,** Spatial plot showing predicted normalized expression of *PDX1* in 8 PCW CosMx representative tissue section (left) after gene expression transfer based on the shared Harmony integrated landscape (See Methods). Spatial plot showing the predicted distribution of M3 mesenchymal population from Ma *et al.*[1] dataset in a 8PCW CosMx representative tissue section (right).

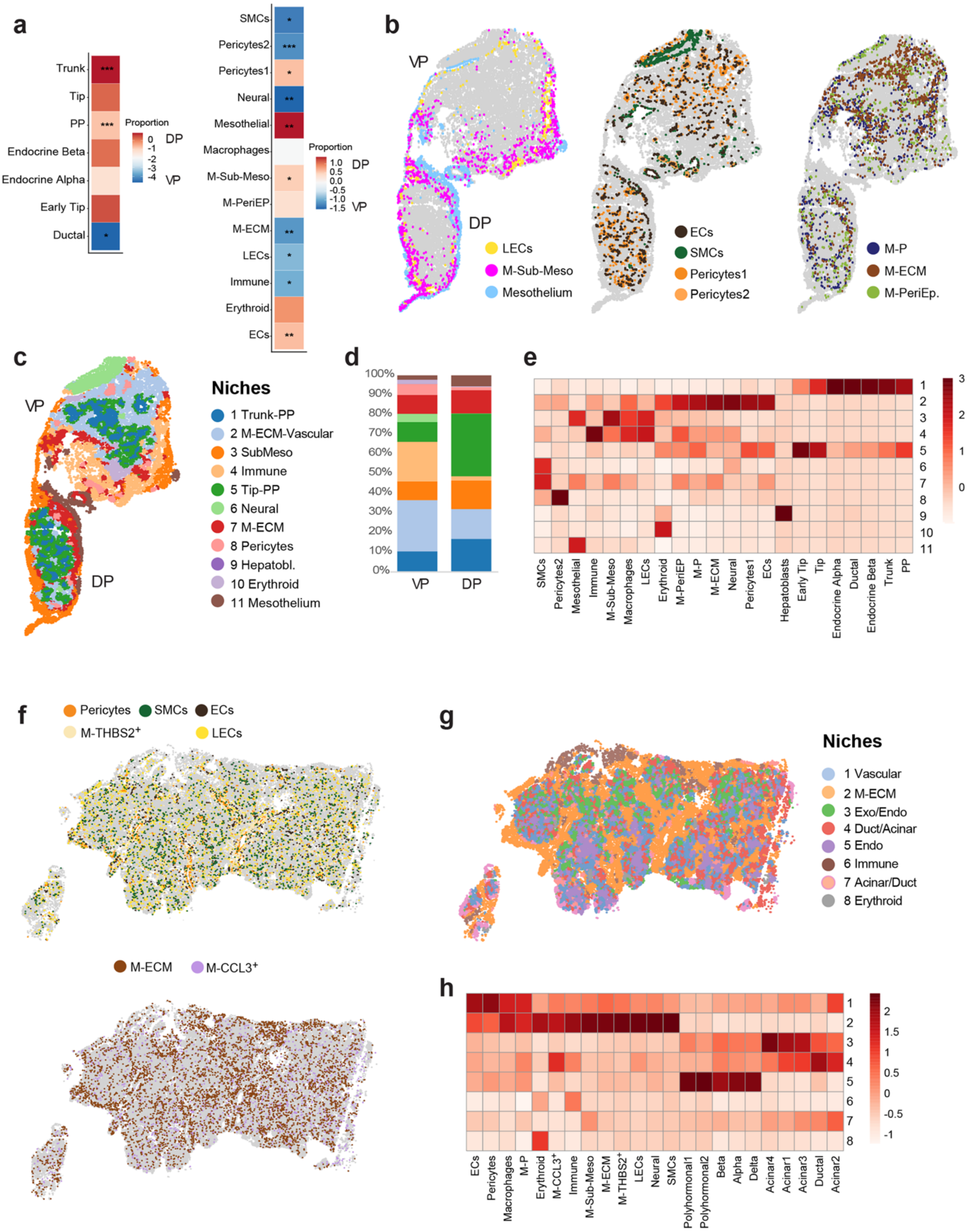


**Extended Data Fig. 4 | Spatial organization of the human foetal pancreas. a,** Heatmaps showing fold changes (FC) in cell-type abundance between dorsal (DP) and ventral (VP) pancreatic buds within the epithelial (left) and non-epithelial (right) compartments. *p<0.05, **p<0.01, ***p<0.001 (Quasi-Likelihood F-Test, adjusted for multiple testing using the Benjamini-Hochberg method). **b,** Spatial distribution of indicated mesenchymal cell populations in a representative 8PCW pancreatic tissue section. **c**, Spatial distribution of Banksy niches in a representative 8PCW pancreatic tissue section. **d**, Banksy niche proportions in DP and VP buds. **e**, Heatmap of cell-type enrichment within 8PCW niches. Colour indicates z-score values. **f,** Spatial distribution of indicated mesenchymal cell populations in a representative 19PCW pancreatic tissue section. **g**, Spatial distribution of Banksy niches in a representative 19PCW pancreatic tissue section. **h**, Heatmap of cell-type enrichment within 19PCW niches. Colour indicates z-score values.

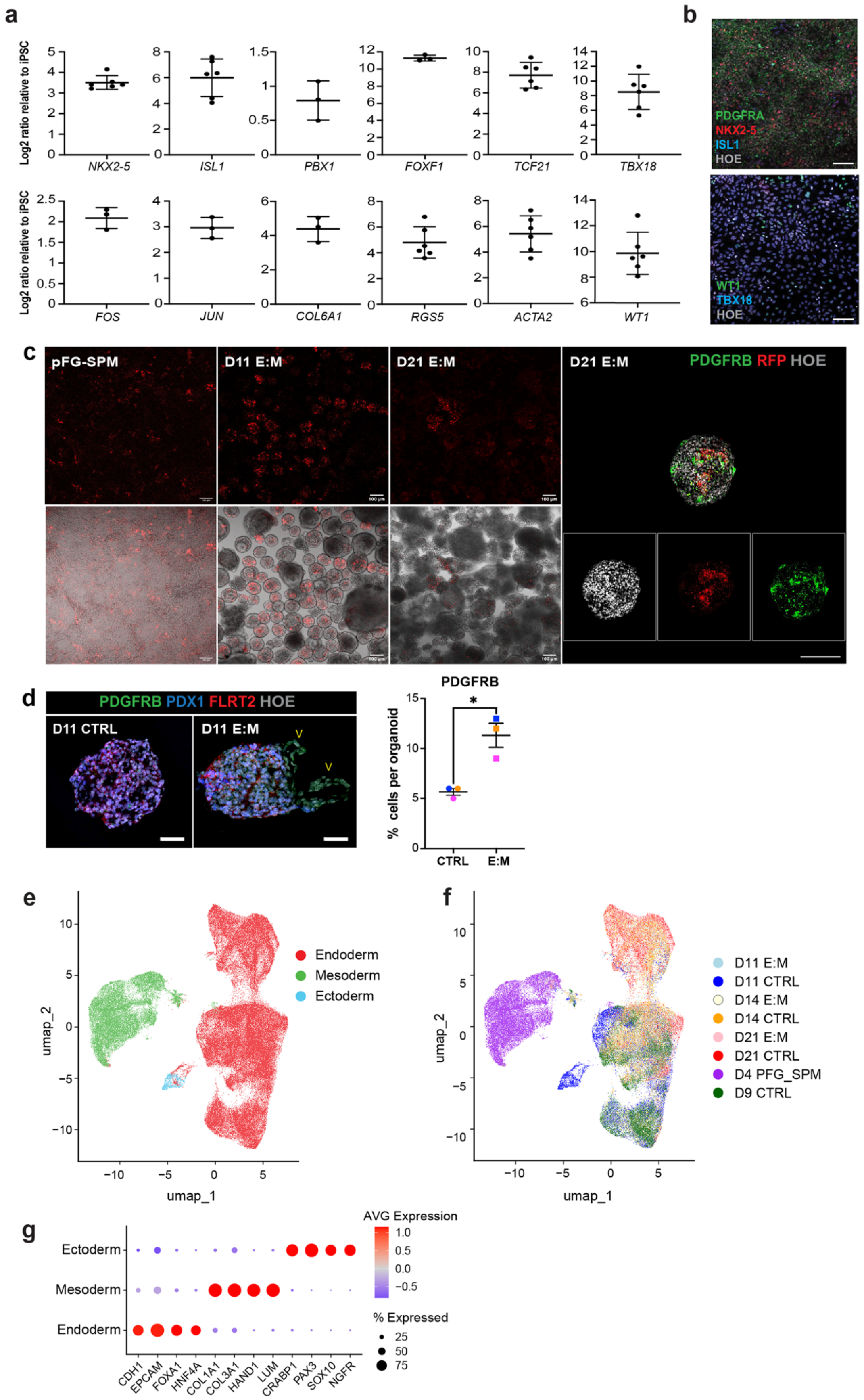


**Extended Data Fig. 5 | Characterization of iPSC-derived posterior foregut splanchnic mesoderm (pFG-SPM). a,** RT-qPCR analysis of indicated mesenchymal genes in iPSC-derived pFG-SPM cells. Data are shown as LOG2 ratio relative to undifferentiated iPSCs. Values shown are mean ± S.D. N=3-6. **b,** Representative immunofluorescence (IF) images of pFG-SPM 2D cultures for the indicated mesodermal and mesothelial markers. Hoechst (Hoe) was used as nuclear counterstaining (grey). Scale bar, 50µm. **c,** Representative native fluorescent images of iPSC-derived pFG-SPM cells transduced with an EF1a/mCherry lentiviral vector before (2D) and after (3D) co-culturing with iPSC-derived PPs at indicated stages. Right panels, representative whole-mount IF images of D21 E:M organoids stained for anti-RFP (red), PDGFRB (green), and Hoechst (grey). Bottom panels show individual fluorescence channels. **d,** Left, representative IF images of D11 control (CTRL) and E:M organoids stained for pancreatic [PDX1 (blue)], mesenchymal markers [PDGFRB (green), FLRT2 (red)] and Hoechst (grey). Arrowheads indicate $PDGFRB^{+}$ cells at the periphery of E:M organoids. Scale bar, 50 µm. Right, plot showing the percentage (%) of $PDGFRB^{+}$ cells relative to the total number of cells per organoid. Off-target spontaneous mesenchymal differentiation observed in CTRL conditions corresponds to approximately 5%. Individual dots represent the mean value of biological replicates. Values shown are mean ± S.E.M. N= 3. (CTRL, n=17; E:M, n=22). *p<0.05 (paired t-test). **e,** UMAP visualization of the integrated multi-lineage organoid sc-RNASeq dataset coloured by main cellular compartment. **f,** UMAP visualization of the integrated multi-lineage organoid sc-RNASeq dataset coloured by sample. **g**, Dot plot showing the expression of selected markers across major compartments. Colour represents scaled AVG expression levels; dot size indicates the % of expressing cells.

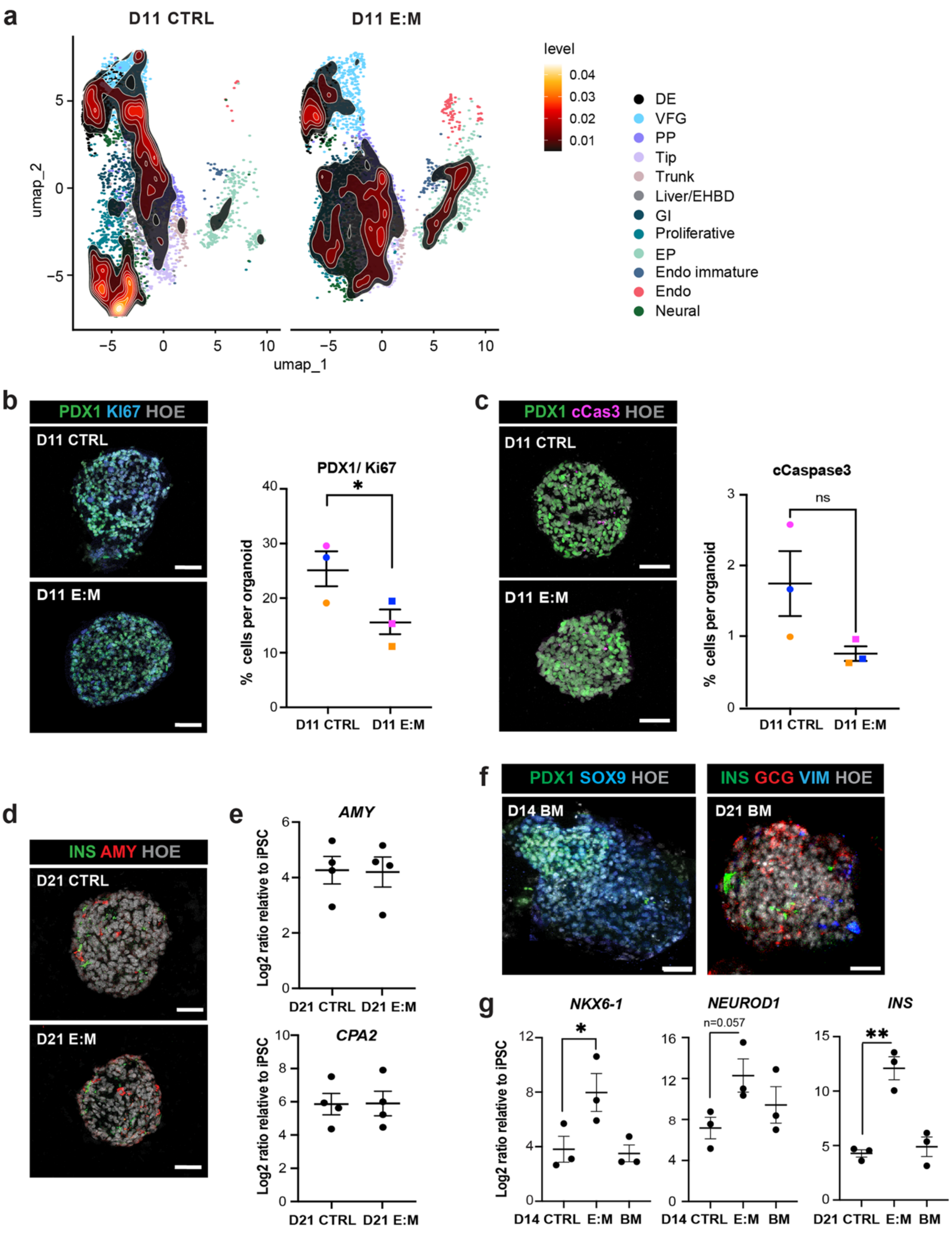


**Extended Data Fig. 6 | iPSC-derived pancreatic mesenchyme promotes a shift in cell-state density towards endocrine differentiation. a,** UMAP and density kernel showing the distribution of cells across clusters from CTRL and E:M organoids. Scale indicates lowest to highest local density. Abbreviations: DE, Definitive Endoderm; EHBD, Extra-Hepatic Biliary

Duct; Endo, Endocrine cells; EP, Endocrine Progenitors; GI, Gastrointestinal; PP, Pancreatic Progenitors; VFG, Ventral Foregut. **b,** Left, representative IF images of D11 CTRL and E:M organoids stained for PDX1 (green), KI67 (blue), and Hoechst (grey). Scale bar, 50µm. Right, scatter plot showing the % of $PDX1^+$ $KI67^+$ cells relative to the total number of cells per organoid. Individual dots represent the mean value of biological replicates. Values shown are mean ± S.E.M. N= 3. CTRL, n=35; E:M, n=41). *p<0.05 (paired t-test). **c,** Left, representative IF images of D11 CTRL and E:M organoids stained for PDX1 (green), Cleaved-Caspase 3 (cCAS3) (magenta), and Hoechst (grey). Scale bar, 50µm. Right, scatter plot showing the % of $cCAS3^+$ cells relative to the total number of cells per organoid. Individual dots represent the mean value of biological replicates. Values shown are mean ± S.E.M. N= 3. (CTRL, n= 19; E:M, n= 10). ns, not significative. **d,** Representative IF images of D21 CTRL and E:M organoids stained for INSULIN (INS) (green), AMYLASE (AMY) (red) and Hoechst (grey). Scale bar, 50µm. **e,** RT-qPCR analysis of selected acinar marker genes (*AMYLASE*, *CPA2)* showing no significant differences between D21 CTRL and D21 E:M organoids. Data are shown as LOG2 ratio relative to undifferentiated iPSCs. Values shown are mean ± S.E.M. N=4. **f,** Representative IF images of D14 and D21 E:M organoids cultured in basal medium (BM), without the supplementation of any pancreatic differentiation cytokines. Scale bar, 50µm. **g,** RT-qPCR analysis of selected endocrine marker genes showing no significant differences between CTRL and E:M organoids cultured in BM conditions. Data are shown as LOG2 ratio relative to undifferentiated iPSCs. Values shown are mean ± S.E.M. N=3. *p<0.05, **p<0.01 (Tukey's multiple comparison after one-way ANOVA).

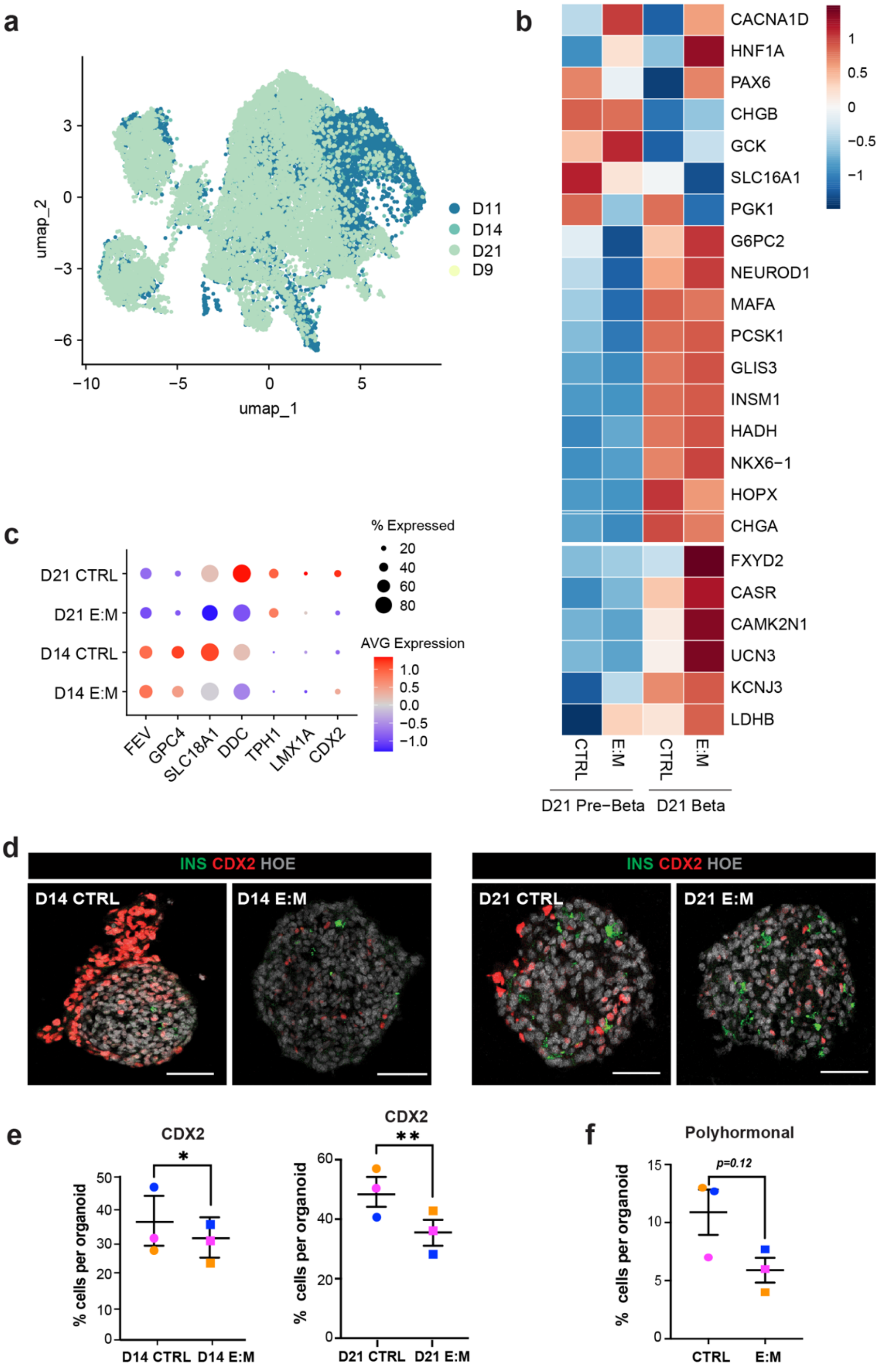


**Extended Data Fig. 7 | Characterisation of endocrine lineage differentiation in E:M pancreatic organoids. a,** UMAP visualization of the endocrine compartment from sc-RNASeq data of CTRL and E:M organoids, coloured by day (D) of differentiation. **b,** Heatmap of scaled expression of indicated genes in beta-cell clusters split per condition. Scale indicates scaled expression levels. **c**, Dot plot showing the expression of selected enterochromaffin marker genes across selected samples. Dot size indicates the % of expressing cells; colour represents scaled AVG expression levels. **d**, Representative IF images of D14 (left) and D21 (right) CTRL and E:M organoids stained for INSULIN (INS) (green), (green), CDX2 (red) and Hoechst (grey). Scale bar, 50μm. **e**, Quantification of $CDX2^{+}$ cells relative to the total number of cells per organoid, shown as %. Individual dots represent the mean value of biological replicates. Values shown are mean ± S.E.M. [Day 14: N= 3 (CTRL, n= 36; E:M, n= 46); Day 21: N= 3 (CTRL, n= 31; E:M, n= 28)]. *p<0.05, ** p<0.01 (paired t-test). **f,** Scatter plot showing the % of polyhormonal ($INS^{+}/GCG^{+}$) cells relative to the total number of cells per organoid at D21. Individual dots represent the mean value of biological replicates. Data are presented as mean ± S.E.M. N= 3. *p=0.12 (paired t-test).

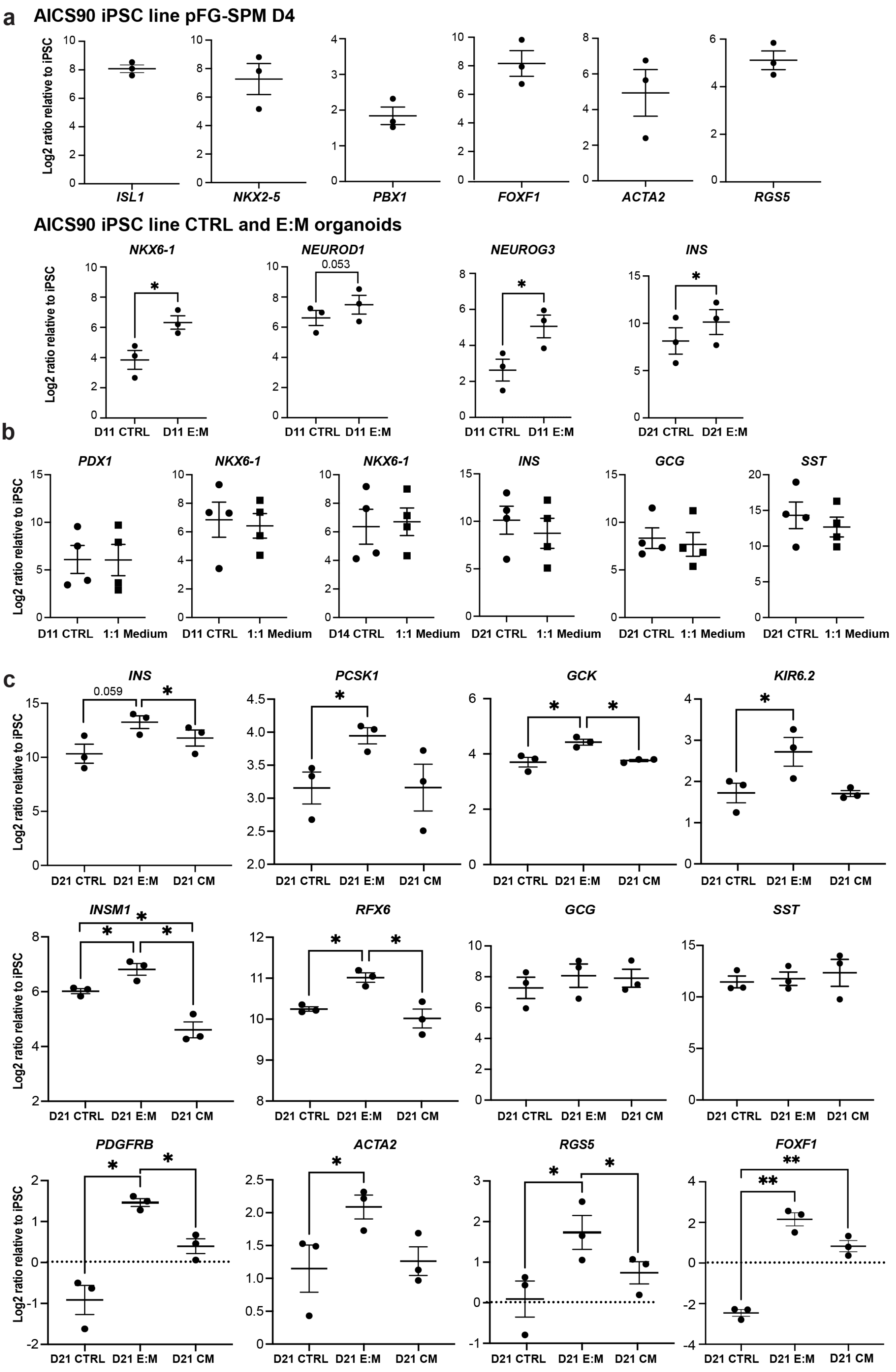


**Extended Data Fig. 8 | E:M multilineage pancreatic organoids from AICS-0090 iPSC line.**

**a,** Characterisation of CTRL and E:M pancreatic organoids from AICS-0090 iPSC line (male donor from the Allen Institute for Cell Science). RT-qPCR analysis showed induction of selected mesodermal and endocrine markers markers in pFG-SPM cells and CTRL and E:M organoids. Data are shown as LOG2 ratio relative to undifferentiated iPSCs. Values shown are mean ± S.E.M. N=3. *p<0.05 (paired t-test). We found that the efficiency in pFG-SPM differentiation and induction of pancreatic endocrine lineage was comparable between the AICS-0090 iPSC line and the HMGUi001-A2 line used throughout the study. **b,** RT-qPCR analysis of indicated markers at indicated days in CTRL organoids cultured in standard differentiation medium or in 1:1 medium used for E:M multilineage organoids (see Methods and Supplementary Information Table S7). Values shown are mean ± S.E.M. N=4. **c,** Comparison of contact-dependent and contact-independent influence of pancreatic mesenchyme in organoids. RT-qPCR analysis of selected beta-cell and mesenchymal marker genes across D21 conditions. Data are shown as LOG2 ratio relative to undifferentiated iPSCs. Values shown are mean ± S.E.M.  N=3. *p<0.05, **p<0.01 (Tukey's multiple-comparison test after one-way ANOVA).

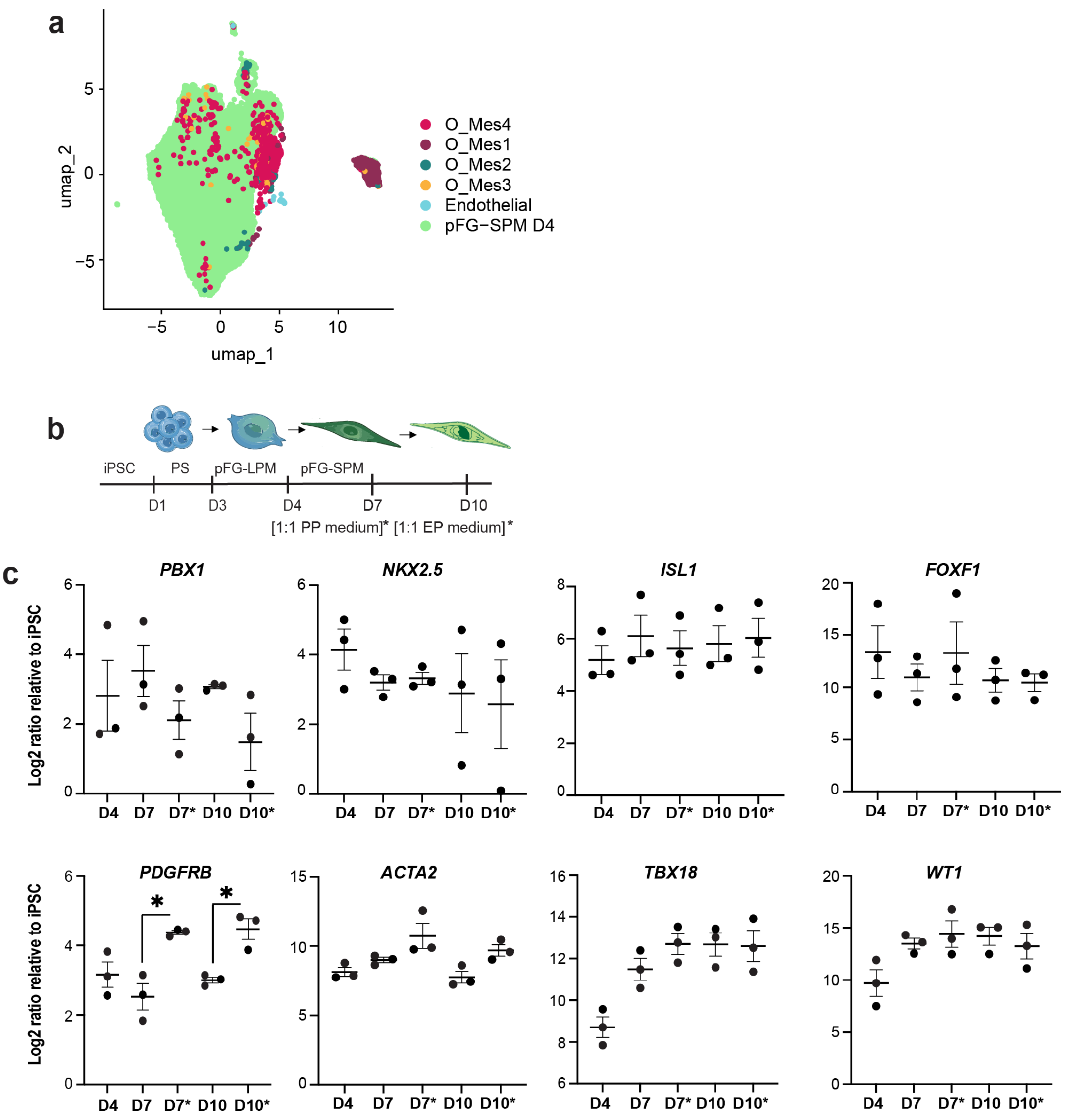


**Extended Data Fig. 9 | Characterization of mesodermal lineage in pancreatic organoid cultures. a,** UMAP visualization of the mesoderm compartment from sc-RNAseq analysis on CTRL and E:M organoids, coloured as in Fig. 5, and Day 4 pFG-SPM cells before coculture. **b,** Schematic representation of the iPSC-derived posterior foregut splanchnic mesoderm (pFG-SPM) culture conditions. pFG-SPM was cultured either in standard mesenchyme medium or exposed to pancreatic differentiation cytokines used for directed differentiation to PP and EP stages (see Supplementary Table S7 for cytokines and differentiation protocols). **c,** RT-qPCR analysis of indicated mesenchymal markers in pFG-SPM cells cultured in control (D7 and D10)

or pancreatic differentiation media (D7* and D10*). Data are shown as LOG2 ratio relative to undifferentiated iPSCs. Values shown are mean ± S.E.M. *$p<0.05$ (paired t-test).

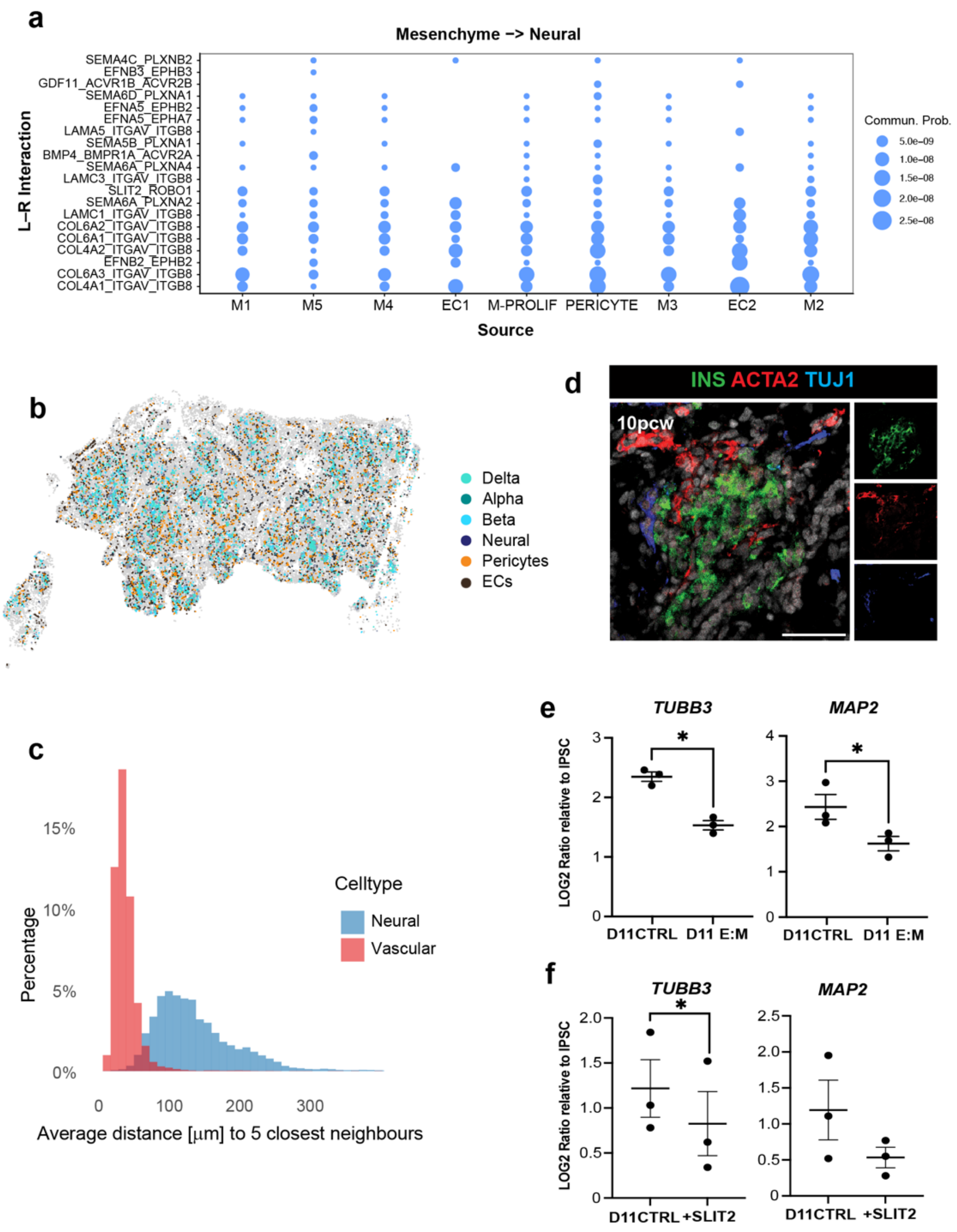


**Extended Data Fig. 10. Characterisation of neural-mesenchymal crosstalk in human foetal development. a,** Conserved ligand-receptor (L:R) interactors between neural cells and indicated mesenchymal and endothelial cell (EC) sources in *in vivo* foetal pancreatic tissue. Analysis was performed using CellChat based on sc-RNASeq data from E:M organoids and publicly available datasets[1]. Dot size represents the calculated communication probability. **b,** Spatial distribution of indicated cell populations in a representative 19PCW pancreatic CosMx section. **c,** Histogram showing the percentage of endocrine cells whose five nearest neural (blue) or vascular (red) cells are within the specified distance. Vascular cells include pericytes and ECs. **d,** Representative IF images of 10 PCW human foetal pancreas showing $INSULIN^{+}$ cells (green), $ACTA2^{+}$ pericytes (red) next to the islet cells, and $TUJ1^{+}$ neural (blue) at the periphery. Scale bars, 50µm. **e,** RT-qPCR analysis of selected neural genes in CTRL, E:M organoids and SLIT2-treated CTRL organoids at D11. Data are shown as LOG2 ratio relative to undifferentiated iPSCs. Values shown are mean ± S.E.M. N=3. *p<0.05 (paired t-test).

### Description of Supplementary Information Tables

**File name: Table S1**

Differentially expressed genes (DEGs) of each population identified in 8PCW CosMx spatial transcriptomic samples. Genes were calculated by comparing each cluster vs. all other clusters with Seurat's FindAllMarkers() function.

**File name: Table S2**

Differentially expressed genes (DEGs) of each population identified in 15PCW CosMx spatial transcriptomic samples. Genes were calculated by comparing each cluster vs. all other clusters with Seurat's FindAllMarkers() function.

**File name: Table S3**

Differentially expressed genes (DEGs) of each population identified in 19PCW CosMx spatial transcriptomic samples. Genes were calculated by comparing each cluster vs. all other clusters with Seurat's FindAllMarkers() function.

**File name: Table S4**

Differentially expressed genes (DEG) of each population identified in sc-RNASeq endoderm compartment clusters. Genes were calculated by comparing each cluster vs. all other clusters with Seurat's FindAllMarkers() function.

**File name: Table S5**

Differentially expressed genes (DEG) of each population identified in sc-RNASeq endocrine compartment clusters. Genes were calculated by comparing each cluster vs. all other clusters with Seurat's FindAllMarkers() function.

**File name: Table S6**

Differentially expressed genes (DEG) of each population identified in sc-RNASeq mesoderm compartment clusters. Genes were calculated by comparing each cluster vs. all other clusters with Seurat's FindAllMarkers() function.

**File name: Table S7**

List of reagents used in the study. **Tab 1:** Primary antibodies. **Tab 2:** Secondary antibodies. **Tab 3:** qPCR primers. **Tab 4:** Reagents and cytokines. **Tab. 5**: Cytokines for pFG_Mesoderm directed differentiation protocol. **Tab 6:** Cytokines for pancreatic directed differentiation, direct co-culture and indirect co-culture protocols.
